# SARS-CoV-2 infection results in lasting and systemic perturbations post recovery

**DOI:** 10.1101/2022.01.18.476786

**Authors:** Justin J. Frere, Randal A. Serafini, Kerri D. Pryce, Marianna Zazhytska, Kohei Oishi, Ilona Golynker, Maryline Panis, Jeffrey Zimering, Shu Horiuchi, Daisy A. Hoagland, Rasmus Møller, Anne Ruiz, Jonathan B. Overdevest, Albana Kodra, Peter D. Canoll, James E. Goldman, Alain C. Borczuk, Vasuretha Chandar, Yaron Bram, Robert Schwartz, Stavros Lomvardas, Venetia Zachariou, Benjamin R. tenOever

**Affiliations:** Department of Microbiology, Icahn School of Medicine at Mount Sinai, New York, NY; Department of Microbiology, New York University, Langone Health, New York, NY; Department of Neuroscience, Icahn School of Medicine at Mount Sinai, New York, NY; Mortimer B. Zuckerman Mind, Brain and Behavior Institute, Columbia University, New York, NY; Department of Neurosurgery, Icahn School of Medicine at Mount Sinai, New York, NY; Department of Otolaryngology- Head and Neck Surgery, Columbia University Irving Medical Center, Vagelos College of Physicians and Surgeons, Columbia University, New York, NY; Department of Pathology and Cell Biology, Columbia University Irving Medical Center, Vagelos College of Physicians and Surgeons, Columbia University, New York, NY; Department of Pathology and Laboratory Medicine, Weill Cornell Medicine, New York, NY; Department of Physiology, Biophysics, and Systems Biology, Weill Cornell Medicine, New York, NY; Division of Gastroenterology and Hepatology, Department of Medicine, Weill Cornell Medicine, New York, NY

## Abstract

SARS-CoV-2 has been found capable of inducing prolonged pathologies collectively referred to as Long-COVID. To better understand this biology, we compared the short- and long-term systemic responses in the golden hamster following either SARS-CoV-2 or influenza A virus (IAV) infection. While SARS-CoV-2 exceeded IAV in its capacity to cause injury to the lung and kidney, the most significant changes were observed in the olfactory bulb (OB) and olfactory epithelium (OE) where inflammation was visible beyond one month post SARS-CoV-2 infection. Despite a lack of detectable virus, OB/OE demonstrated microglial and T cell activation, proinflammatory cytokine production, and interferon responses that correlated with behavioral changes. These findings could be corroborated through sequencing of individuals who recovered from COVID-19, as sustained inflammation in OB/OE tissue remained evident months beyond disease resolution. These data highlight a molecular mechanism for persistent COVID-19 symptomology and characterize a small animal model to develop future therapeutics.

## INTRODUCTION

SARS-CoV-2 is a respiratory RNA virus that emerged in 2019 and is associated with a variety of clinical phenotypes ranging from asymptomatic to more severe disease generally referred to as coronavirus-induced disease (COVID-19)(Yuki et al., 2020). In most cases among young and healthy individuals, COVID-19 is characterized by a mild flu-like illness and includes limited respiratory tract congestion, fever, myalgia, headache, and anosmia (Hu et al., 2021; Wiersinga et al., 2020; Zhu et al., 2020). Amongst older populations, especially males and those with co-morbidities, COVID-19 can result in severe respiratory distress, multi-organ complications, and death (Bhopal and Bhopal, 2020; Yuki *et al*., 2020).

Regardless of age or underlying health, virus infection is thought to impair host transcriptional and translational processes to enhance replication (Banerjee et al., 2020; Schubert et al., 2020; Thoms et al., 2020). As a result, infected cells are unable to elicit a Type I interferon (IFN-I) response, a central mediator for initiating the host’s antiviral defenses through the upregulation of hundreds of antiviral interferon-stimulated genes (ISGs) (Blanco-Melo et al., 2020; Hadjadj et al., 2020). During a SARS-CoV-2 infection, induction of IFN-I largely derives from uninfected cells such as resident macrophages and other phagocytic cells (Grant et al., 2021). Interestingly, despite blocking many aspects of the host antiviral response, SARS-CoV-2 infection results in persistent signaling of the NFκB transcription factor family, culminating in high levels of proinflammatory cytokines (i.e. interleukin 6, IL6) and chemokines (i.e. CXCL10) (Blanco-Melo *et al*., 2020; Lucas et al., 2020; Nilsson-Payant et al., 2021). As a result of these dynamics, high levels of neutrophils and monocytes amass in the respiratory tract as the virus propagates in an environment with minimal antiviral defense engagement - further exacerbating the inflammatory environment. Virus infection results in extensive damage to the bronchial epithelium and pulmonary edema due to the proinflammatory cytokine response and/or a loss of normal lung function (Hu *et al*., 2021; Wiersinga *et al*., 2020; Zhu *et al*., 2020).

Characterization of SARS-CoV-2 biology has identified ACE2 and a small subset of proteases that enable viral entry (Cantuti-Castelvetri et al., 2020; Daly et al., 2020; Hoffmann et al., 2020). Despite the expression of these factors on multiple tissues, productive SARS-CoV-2 infection appears to be largely contained in the respiratory tract (Hu *et al*., 2021; Wiersinga *et al*., 2020; Zhu *et al*., 2020). Selective localization in the airways however is not a product of viral tropism or receptor expression but rather a result of the systemic IFN-I response which radiates from the site of infection. For example, human organoid models have demonstrated productive infection of diverse tissues *ex vivo* despite being rarely observed *in vivo* (Asano et al., 2021; Lamers et al., 2020; Yang et al., 2020). This phenomenon can also be modeled in the golden hamster, arguably the best small animal model for COVID-19, which demonstrates consistent infection of the respiratory tract and olfactory epithelium with only sporadic isolation of virus from other tissues unless IFN-I biology is disrupted (Boudewijns et al., 2020; Hoagland et al., 2021; Imai et al., 2020; Sia et al., 2020; Zazhytska et al., 2021). This same phenomenon can be observed when infected individuals are immunosuppressed (Asano *et al*., 2021; Bastard et al., 2020; Puelles et al., 2020). While it remains unclear how common infection of distal tissues is during a SARS-CoV-2 infection, system-wide inflammation is consistent (Hu *et al*., 2021; Wiersinga *et al*., 2020; Zhu *et al*., 2020). Together these data suggest that the molecular underpinnings of acute COVID-19 are a by-product of the damage caused by the virus and the systemic response that ensues.

In most individuals, virus infection is successfully cleared with the appearance of neutralizing antibodies to the Spike (S) attachment protein. Generally, the appearance of the humoral response correlates to resolution of the symptoms associated with SARS-CoV-2 (Bartsch et al., 2021; Brouwer et al., 2020; Garcia-Beltran et al., 2021). However, a growing body of evidence suggests that in a subset of individuals, SARS-CoV-2 infection results in prolonged complications including shortness of breath, persistent fevers, fatigue, depression, anxiety, and a state of chronic impairment of memory and concentration known colloquially as “brain fog”. The direct cause of these impairments, known collectively as “long COVID” or post acute sequalae of COVID-19 (PASC), is currently unknown (Nalbandian et al., 2021; Sudre et al., 2021).

To better understand the prolonged effects caused by SARS-CoV-2 infection, we focused on the golden hamster as a model system. The hamster model has proven to largely phenocopy COVID-19 biology without any requirement for SARS-CoV-2 adaptation and has demonstrated a propensity to display severe lung morphology and a tropism that matches what is observed in human patients(de Melo et al., 2021; Hoagland *et al*., 2021; Imai *et al*., 2020; Sia *et al*., 2020). Here we show that while both IAV and SARS-CoV-2 induce a systemic antiviral response, only the latter infection results in sustained inflammatory pathology that extends well beyond clearance of the primary infection. As this sustained inflammation also correlates with behavioral abnormalities, we propose that this biology may underlie prolonged symptomology that results from SARS-CoV-2 infection.

## RESULTS

### SARS-CoV-2- and IAV-infected hamsters induce a host response that mirrors human biology and resolves within two weeks post infection

To define the unique characteristics of SARS-CoV-2 infection that may contribute to persistent symptomology, we performed a longitudinal study in hamsters infected with either SARS-CoV-2 or Influenza A virus (IAV). These data demonstrated that both respiratory RNA viruses could replicate in the lungs of the golden hamster, albeit with different rates of clearance, consistent with what has been reported elsewhere (Figure 1A-B, S1A-B)(Hoagland *et al*., 2021). IAV challenge resulted in peak titers of 10^7 plaque forming units per gram of lung tissue (pfu/g) on day three followed by a sharp decline in infectious material – showing a complete loss of infectivity by day seven post-infection (Figure 1A). For SARS-CoV-2, we also observed peak viral titers on 3 days post-infection (dpi) (∼10^8pfu/g), however these levels persisted till day 5 before declining (Figure 1B). Despite the difference in controlling overall virus levels, in both model systems, no infectious virus could be isolated on day seven although trace levels of RNA remained detectable via quantitative reverse-transcription-based PCR (qRT-PCR) for the nucleoprotein of influenza (NP) as well as the sub-genomic mRNA of nucleocapsid (N) from SARS-CoV-2 (Figure 1A-B and S1A-B). Based on these data, we decided to focus on day three to compare the acute host response to these two respiratory infections.

**Figure 1.**
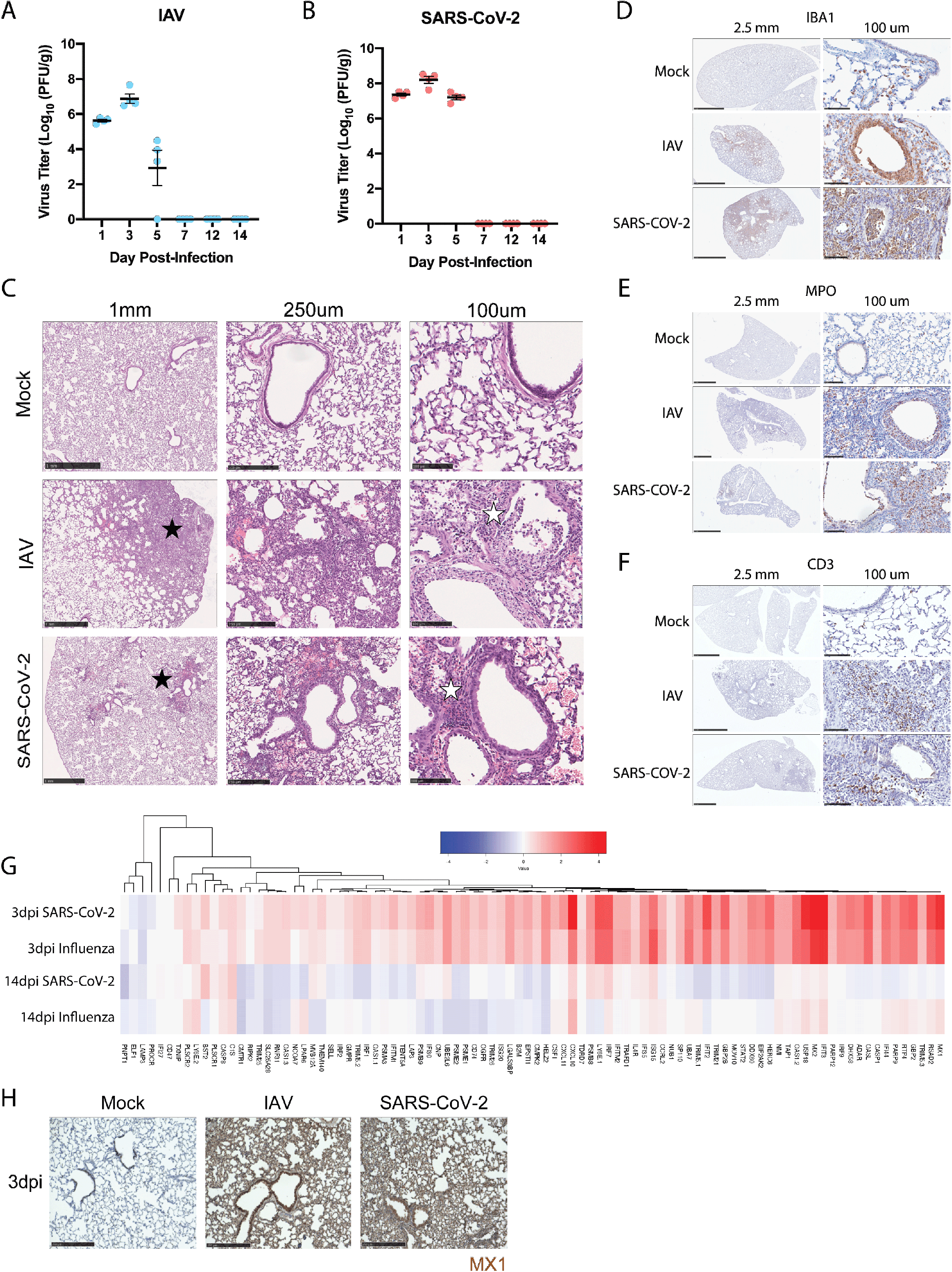
SARS-CoV-2 and IAV infections induce clinically representative lung pathology and are cleared by 14dpi in the hamster model of disease. **(A-B)** Titer data computed as plaque forming units per gram (PFU/g) of lung from hamsters infected with IAV (A/California/02/2009) or SARS-CoV-2 (USA/WA1/2020) on days indicated. (**C**) H&E staining of hamster lungs treated with PBS (Mock), IAV, or SARS-CoV-2 at 3dpi. Histological analysis at various magnifications denoting infiltration (black star) or bronchiolar epithelial necrosis (white star). (**D-F**) Immunohistochemical labeling for (D) IBA1, (E) MPO, and (F) CD3, were used to label macrophage, neutrophil, and T cell populations, respectively, in the lungs of mock-, IAV-, and SARS-CoV-2-infected hamsters at 3dpi. Size of inset scale bars matches length described in column headers. **(G)** RNA-sequencing of lungs from SARS-CoV-2- and IAV-infected hamsters was evaluated at 3dpi and 14dpi. Heatmap depicting log2 fold-change of IFN-I response genes (derived from HALLMARK_INTERFERON_ALPHA_RESPONSE gene set) compared to mock-infected animals was generated for these groups. **(H)** Immunohistochemical labeling for interferon stimulated gene MX1 was assessed in lungs of mock-infected, IAV-infected, or SARS-CoV-2-infected hamsters at 3dpi.

To compare the pathology resulting from IAV vs. SARS-CoV-2, we examined cross sections of the hamster lung at 3dpi using various histological techniques that were evaluated by a board-certified pathologist (Figure 1C-F). Hematoxylin and Eosin (H&E) staining on the lungs of hamsters infected with either IAV or SARS-CoV-2 revealed large areas of intense staining (Figure 1C, black stars) that, at higher magnification, were shown to be comprised of nuclei suggesting hypercellularity and the infiltration of inflammatory cells into both alveolar compartments and bronchiolar airway spaces (white stars). To better characterize the cellular content of the pulmonary inflammatory infiltrate, immunohistochemical (IHC) staining was used to label macrophages (IBA1) (Figure 1D), neutrophils (MPO) (Figure 1E), and T cells (CD3) (Figure 1F) on these same cross-sections. These efforts demonstrated that intensely hematoxylin-stained regions of the lung sections with either virus showed high levels of positivity for all three cell types, with neutrophils and macrophages predominating (Figures 1C-F). One notable difference between these virus models was that SARS-CoV-2 induced pulmonary infiltration, centrally located around bronchioles and larger airway structures (Figure 1C, black star). In contrast to the lung, examination of distal tissues including kidney and heart showed moderate to no pathological features at 3dpi for either virus (Figure S1C-D). We observed no signs of cellular infiltration in the kidney at this time point, whereas in the heart, an organ often associated with COVID-19 complications (Satterfield et al., 2021), we noted some evidence for inflammation and leukocytic infiltrate in response to both viruses (Figure S1C-D, green stars). Together, these data suggest that the hamster phenocopies many of the histological characteristics seen in the human response to IAV or SARS-CoV-2 during acute infection.

To characterize the molecular dynamics of these model systems, we next performed RNA-Seq on infected lung from 3dpi (Figures 1G and S1E-F). These data identified ∼100 differentially expressed genes (DEGs) with a P-adjusted value of less than 0.1 in both SARS-CoV-2- and IAV-infected lungs. Gene set enrichment analyses (GSEA) against hallmark gene ontology sets implicated activation of the IFN-I and IFN-II response as well as TNFα and IL2 signaling in response to either infection (Figure S1G). These data could be further corroborated by immunohistochemistry of fixed lung tissue probed for the ISG MX1 (Figure 1H). Together, these data suggest that the host response to IAV and SARS-CoV-2 in the lung, kidney, and heart is comparable with the only major difference being that titers of SARS-CoV-2 were maintained through 5dpi.

We next examined the transcriptional response to SARS-CoV-2- and IAV-infected hamsters at 14dpi, ∼one week post clearance (Figure 1A-B, G). In contrast to the inflammation observed at 3dpi, sequencing SARS-CoV-2- or IAV-infected lung tissue at 14dpi showed minimal signs of an antiviral response (Figure 1G). Together, these data demonstrate that the golden hamster model shows a robust acute response in the respiratory tract that successfully resolves both IAV or SARS-CoV-2 infection.

### Transcriptional profiling of peripheral organs during active or resolved IAV vs. SARS-CoV-2 infections

To corroborate the clinical validity of the SARS-CoV-2 acute hamster data, we compared our RNA-seq analyses to published results from lungs of COVID-19 deceased individuals that still had high viral loads at the time of death (Figure 2A-C) (Butler et al., 2021). In agreement with the published data, we find transcriptional signatures from both groups were dominated by a marked upregulation of the IFN-I response as well as TNFα signaling via NFκB (Blanco-Melo *et al*., 2020; Hoagland *et al*., 2021; Nilsson-Payant *et al*., 2021).

**Figure 2.**
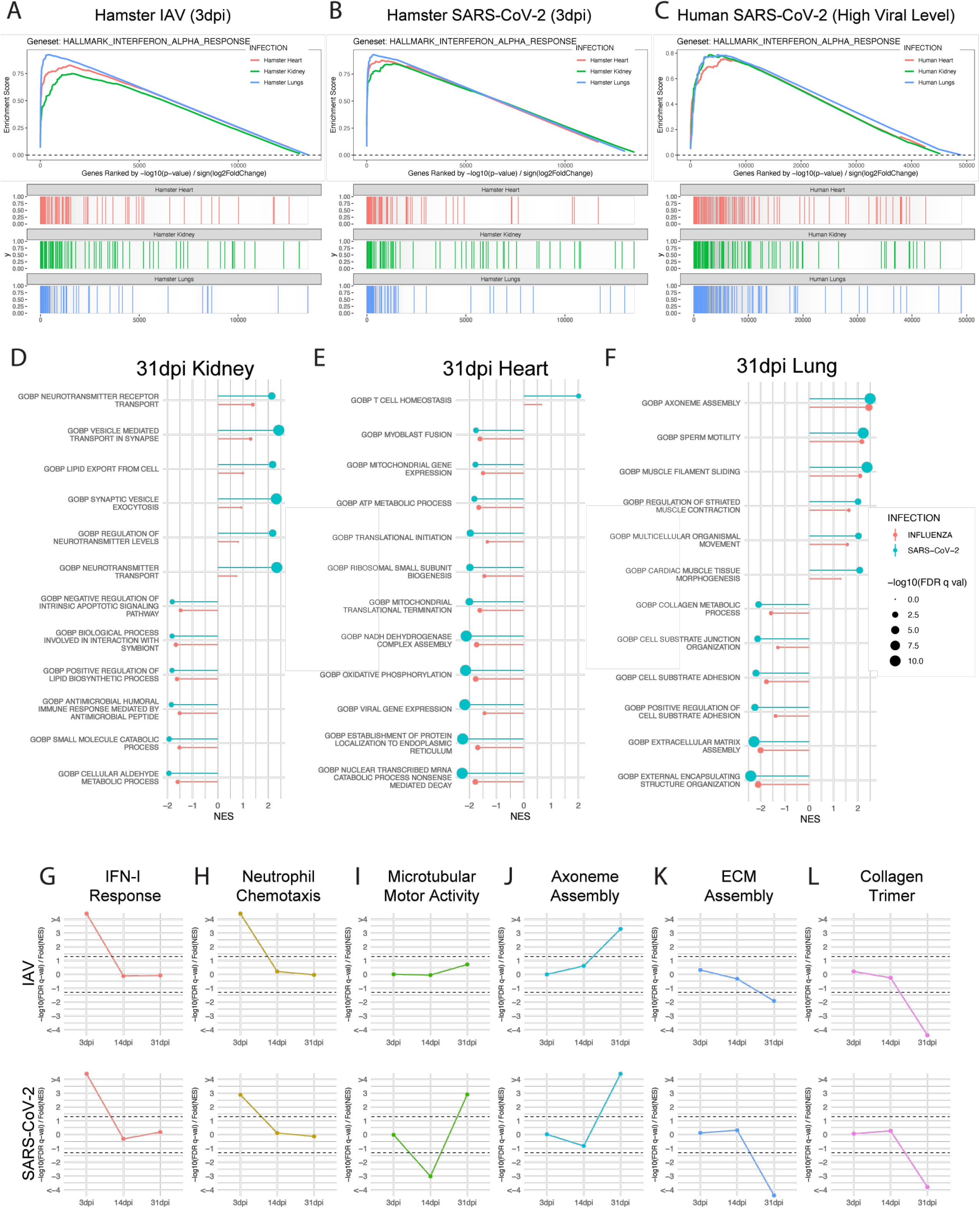
Transcriptional profiling of peripheral organs during active or resolved IAV vs. SARS-CoV-2 infections. **(A-C)** Lungs (blue), kidneys (green), and hearts (red) of SARS-CoV-2-, IAV-, and mock-treated hamsters were harvested at 3dpi and transcriptionally profiled via RNA-seq. Differential expression analysis was conducted between infected and mock-infected groups with DESeq2 and analyzed via Gene Set Enrichment Analysis (GSEA) for enrichment of indicated gene sets. Enrichment analysis results for all three tissue types are displayed in a GSEA enrichment plot for **(A)** IAV vs. Mock and **(B)** SARS-CoV-2 vs. Mock comparisons. **(C)** Similar transcriptomic analyses were conducted on RNA-seq data generated from human lung, heart, and kidney samples obtained from the post-mortem tissues of COVID-19-infected and control donors. Results from enrichment analyses are shown as a GSEA enrichment plot. **(D-F)** Differential expression analysis of RNA-Seq data derived from **(D)** Lungs, **(E)** kidneys, and **(F)** hearts of SARS-CoV-2-, IAV-, and mock-infected hamsters at 31dpi. Differential expression results were assessed via GSEA to test for enrichment of gene sets present in the MSigDB C5 gene set collection, which contains curated gene sets derived from the Gene Ontology resource. Significant ontological enrichments for SARS-CoV-2 vs. Mock differential expression analysis were further processed via REVIGO to remove redundant enrichments. The highest ranked non-redundant positive and negative enrichments for each organ are plotted by their normalized enrichment score (NES) (line magnitude) and significance (-log10(FDR q-value)) (dot size). GSEA enrichment for these same gene sets in IAV vs. Mock differential expression data for the same tissue are plotted side-by-side for comparison. **(G-L)** GSEA analysis from lung sequencing data from IAV-, SARS-CoV-2, and mock-infected hamster lungs at 3, 14, and 31dpi using curated gene ontology and human phenotype ontology gene sets. Directional significance of enrichment was plotted over time for (**G**) IFN-I response (GOBP_RESPONSE_TO_TYPE_I_INTERFERON) (**H)** neutrophil chemotaxis (GOBP_NEUTROPHIL_CHEMOTAXIS) (**I**) microtubular motor activity (GOMF_ATP_DEPENDENT_MICROTUBULE_MOTOR_ACTIVITY) (**J**) axoneme assembly (GOBP_AXONEME_ASSEMBLY) (**K**) extracellular matrix (ECM) assembly (GOBP_EXTRACELLULAR_MATRIX_ASSEMBLY) (**L**) and collagen trimer-associated genes (GOCC_COLLAGEN_TRIMER). Dotted lines show the calculated statistic for FDR q-val = 0.05 for positive and negative enrichment; thus, any points falling outside the dotted lines have FDR q-val of < 0.05.

As the host response to acute IAV vs. SARS-CoV-2 were comparable in the respiratory tract, we next sought to characterize distal tissues and expand our characterization to time points representing both active, as well as resolved infections. While COVID-19 symptoms usually resolve within four weeks post infection onset, symptoms can persist significantly longer in a subset of patients. Patients demonstrating symptoms lasting longer that four weeks following infection have now been clinically defined as having long COVID or PASC (Aiyegbusi et al., 2021). To this end, we conducted additional transcriptional profiling on the lung (blue), heart (red), and kidneys (green) from hamsters infected with SARS-CoV-2 or IAV at 3dpi and 31dpi – a timepoint where any symptom-generating pathology would be clinically defined as long COVID in a human patient (Figures 2A-B and S2A-C). Moreover, these data were cross-referenced to matching tissues derived from human COVID-19 cadavers actively infected at the time of death (Figure 2C). These comparisons encompassed more than 50 samples at both early and late time points which clustered based on tissues from which they derived (Figure S2D).

To first assess the acute response in a more systemic fashion, we utilized GSEA to characterize curated ontology gene sets from the aforementioned tissues. These efforts implicated a strong acute induction of the IFN-I response (FDR q-val < 0.0001) in all three organs following either SARS-CoV-2 or IAV infection, that were also evident in corresponding human tissues (Figure 2A-C, S2E-G). Also of note, IFN-I signatures in the lung of hamsters and COVID-19 cadavers were also accompanied by upregulation of IFN-I-associated pathways including NFκB- and IL6-associated target genes (Figure S2H-J). Other enriched pathways induced when directly comparing SARS-CoV-2 to IAV infection included positive regulation of complement activation in the kidney and negative regulation of calcium channel formation in the heart, although these enrichments were relatively minor in comparison to the IFN-I signatures (Figure S2E-H). Together these data corroborate earlier studies and provide further support for the use of the golden hamster as a model for acute SARS-CoV-2 pathology (Hoagland *et al*., 2021; Imai *et al*., 2020).

Having established the hamster as a clinical proxy for systemic acute pathology in response to a respiratory infection, we next sought to extrapolate these findings to any possible long-term consequences. To this end we performed similar analysis on lung, heart, and kidney tissues from 31dpi, representing a time point greater than two weeks past disease resolution of the lungs (Figure 1A-B, 2D-F). In agreement with the clearance of virus, these analyses failed to show any significant enrichment of IFN-I-or chemokine related signatures in any of the tissues examined (Figure 2D-F). Instead, tissue-specific annotations identified various biologies involved in kidney resorption capacities and heart metabolism (Figure 2D-E). In the lung, GSEA at 31dpi implicated general pathways of repair and regeneration (Figure 2F). Amongst these was the biogenesis of cilia and airway repair in the lung following infection which drives the upregulation of genes involved in axoneme assembly and filament sliding, which are also involved in pathways that identified sperm motility (Figure 2F).

To further visualize the development of the respiratory ontologies over time, we combined all of the sequencing performed on days 3, 14, and 31dpi and mapped their significance (Figure 2G-L). These data corroborated our earlier findings that SARS-CoV-2 and IAV infections were resolved by day 14 as the IFN-I response and neutrophil chemotaxis showed a lack of significant enrichment at both days 14 and 31 post-infection despite their strong induction at 3dpi (Figure 2G-H). Next, we assessed microtubular motor activity and ciliary assembly ontologies over this longitudinal comparison to better understand the dynamics of bronchiolar repair given this biology was enriched in both viral infection models at 31dpi (Figure 2F). While neither infection cohort showed an enrichment in ciliary-related ontologies at 3dpi, there appeared to be a disparity between the two groups at 14dpi (Figure 2I-J). At this time point, SARS-CoV-2-infected lungs uniquely displayed a significant negative enrichment of microtubular motor activity and a trending negative enrichment of axoneme assembly ontologies. As ciliary loss is part of the acute lung pathology following respiratory virus infection (Hoagland *et al*., 2021), we found the decline of microtubules and axoneme assembly-related genes specifically in response to SARS-CoV-2 noteworthy. These findings suggest that SARS-CoV-2-induced transcriptional aberrations may still be prevalent past 14dpi, even in the absence of infectious virus or proinflammatory transcriptional profiles. These data suggest that SARS-CoV-2 induced damage may be more severe and/or persist for a longer duration in the respiratory tract as compared to IAV. However, by 31dpi, the significant increase in microtubular motor activity and axoneme assembly likely reflect active regeneration of the ciliary machinery. This biology also corresponds with the downregulation of genes associated with extracellular matrix assembly and collagen-trimer-related genes which are involved in tissue regeneration (Figure 2K-L)(Darby and Hewitson, 2007; Hogan et al., 2014; Stone et al., 2016). The shared trends observed for the GSEAs suggests a resolving repair response at 31dpi following either IAV or SARS-CoV-2 infection.

### Histological characterization of lung, heart, and kidney tissue in response to SARS-CoV-2 or IAV 31dpi

To assess long term organ damage independent of the transcriptional response, we profiled lung, heart, and kidney by histological analyses following IAV or SARS-CoV-2 infection at 31dpi (Figure 3 and S3). H&E staining revealed that both SARS-CoV-2- and IAV-infected lungs maintained their general structure but displayed numerous abnormalities. Most prominent amongst these pathologies was lambertosis (also known as peribronchiolar metaplasia) (black stars), a clinical finding in which alveolar epithelial cells undergo metaplastic transformation to become bronchiolar-epithelium-like in appearance (Figure 3A). This process generally occurs in response to severe respiratory trauma and can result in functional respiratory defects (Allen, 2010; Taylor et al., 1992; Wright et al., 2020). Notably, while lambertosis was generally localized to alveolar clusters immediately surrounding bronchioles, in the SARS-CoV-2-infected lungs, peribronchiolar metaplasia could be seen expanding across large areas well beyond the general vicinity of the bronchiole, thus visible even at minimal magnification as a more intensely stained area (white star) (Figure 3A). Furthermore, lungs infected by both viruses showed signs of enlarged airway spaces and residual inflammation characterized by monocytes and neutrophils visible in the alveolar spaces (red stars) (Figure 3A). This residual inflammation is in agreement with our transcriptional profiling data which found *Cd177* and *Ly6d*, neutrophil- and monocyte-associated genes, respectively, were significantly upregulated in lungs infected by SARS-CoV-2 at 31dpi (Figure S2A). To confirm these findings, IHC staining was performed to label macrophage (IBA1), neutrophil (MPO), or T-cell (CD3) populations in histological sections of lungs of infected hamsters at 31dpi (Figure S3A-C). In contrast to lungs from mock-infected animals, lungs from IAV- and SARS-CoV-2-infected animals showed localized areas of hypercellularity that stained strongly positive for both neutrophil and macrophage populations (Figure S3A-C). Intriguingly, these hypercellular areas oftentimes appeared co-localized with areas of lambertosis, which could be distinguished via thickened alveolar walls compared to surrounding healthy and mock alveolar tissues. Additionally, in line with our sequencing, which identified a moderate and resolving repair response at 31dpi, Verhoeff Van Gieson staining, which labels collagen and elastin fibers, showed no obvious signs of fibrotic activity, collagen deposition, or elastin degradation in response to either infection (Figure S3D).

**Figure 3.**
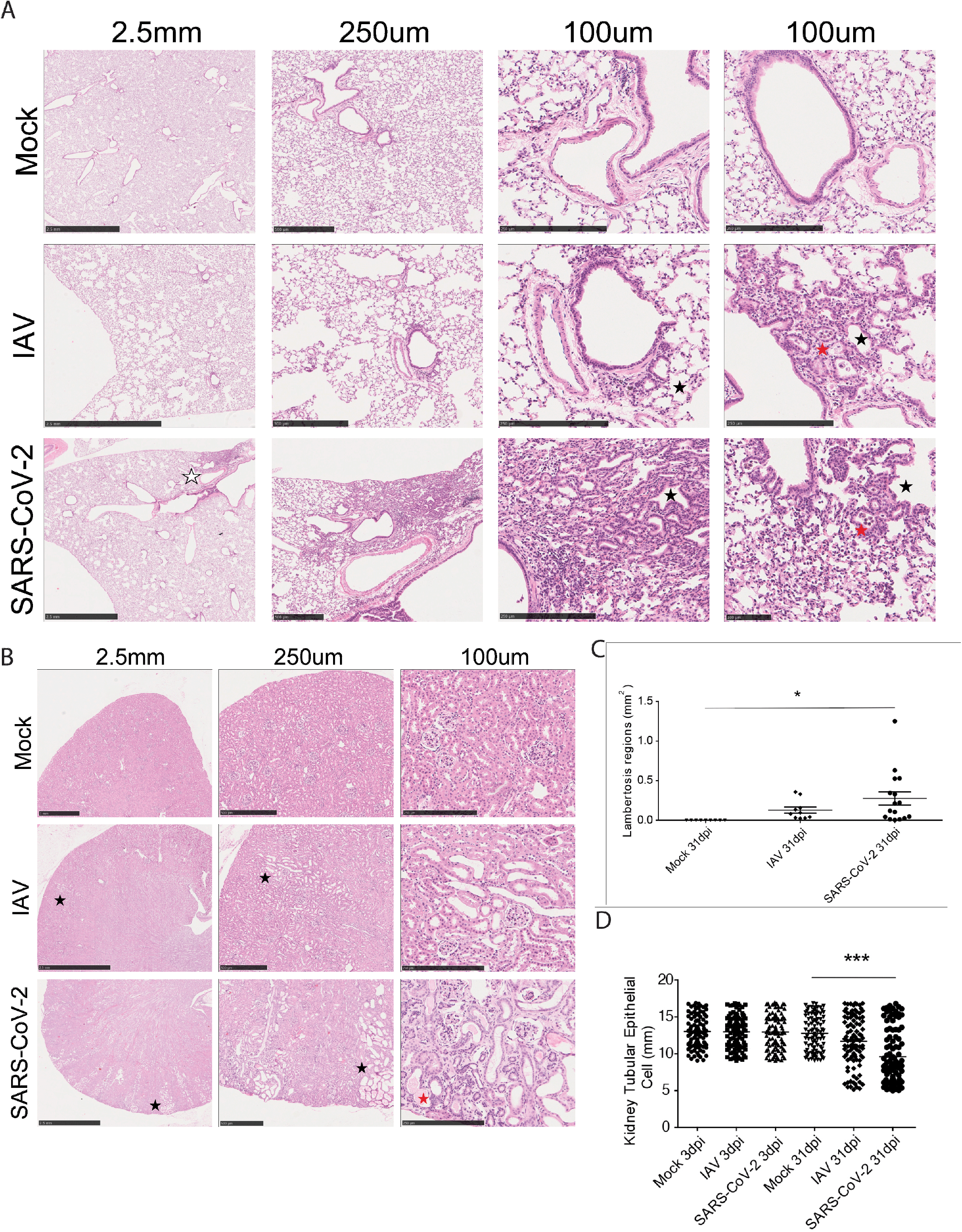
Morphological characterization of lung, heart, and kidney in response to SARS-CoV-2 or IAV at 31dpi. **(A)** H&E staining on lungs of SARS-CoV-2-, IAV-, and mock-infected hamsters at 31dpi. Histological analysis of lungs highlighting lambertosis at both low magnification (white stars) and at higher magnification (black stars) as well as residual immune infiltration into lung parenchyma (red stars). **(B)** H&E staining on kidneys collected from the same infection groups as (A). Black and red stars denote areas of tubular atrophy and sproteinaceous fluid buildup, respectively. **(C-D) Quantification of** lambertosis (**C**) and average tubular epithelial size (**D**) were quantified via morphometric image analysis. Error bars display standard error mean, and significance was quantified via one-way ANOVA with Tukey’s Multiple Comparison Test (*p<0.05, **p<0.01, ***p<0.001)

Given the persistent gene signatures on day 31 post-SARS-CoV-2 infection and the histological changes observed in the lung, we next assessed two distal organs via H&E staining: the kidneys and heart (Figure 3B and S3D). In the heart we observed complete resolution of leukocytic infiltration at 31dpi with no noteworthy histological signatures in response to infection (Figure S3E). In the kidney however, SARS-CoV-2-infected animals displayed areas of tubular atrophy characterized by thinning of tubular cells and widening of the tubular lumen (black stars) (Figure 3B). Closer examination also revealed the presence of proteinaceous fluid in the interstitial space surrounding these tissues (red stars). Examination of the kidneys at this time point from IAV-infected hamsters showed similar pathological findings (black stars); however, the affected areas appeared smaller and less numerous than in SARS-CoV-2-infected hamsters consistent with the notion that IAV-induced damage is less severe than that of SARS-CoV-2 in this small animal model.

To better assess the extent of infection-induced scarring, we performed quantitative morphometric analyses on these histological images. Quantification of lambertosis and airway size showed that these pathologies were indeed significantly greater in the lungs of SARS-CoV-2-infected animals (Figure 3C and S3F). An identical trend was also clearly visible with respect to tubular atrophy and SARS-CoV-2 (Figure 3D). Together, these data demonstrate that both SARS-CoV-2 and IAV infections present similar histological signatures in the lungs and in other peripheral organs. However, despite comparable host responses, we do note a greater severity of scarring in SARS-CoV-2 infection, which, given its nature, may predispose infected individuals to greater functional defects in the affected organs.

### SARS-CoV-2 induces unique neural transcriptional profiles compared to IAV

Given that long COVID may also involve neurological and neuropsychiatric symptomology (Sudre *et al*., 2021), we next assessed the consequences of SARS-CoV-2 infection on the nervous system. For these studies, we transcriptionally profiled several areas of the nervous system from 3 and 31dpi cohorts. More specifically, the areas surveyed included the olfactory bulbs, medial prefrontal cortex (mPFC), striatum, thalamus, cerebellum, and trigeminal ganglion (tissues collected as depicted in Figure 4A). These areas were chosen either due to their previously documented positivity for SARS-CoV-2 transcripts in human patients (olfactory bulb, trigeminal ganglion) or due to their functional importance in sensory, motor, cognitive, and/or affective processes—all of which have been noted to be altered in subsets of long COVID patients (Carrera and Bogousslavsky, 2006; Cox and Witten, 2019; Euston et al., 2012; Gheusi et al., 2000; Thalakoti et al., 2007). Matched tissues from hamsters infected with IAV were also collected for comparison. Following tissue processing, brain regions from 3dpi were surveyed for the presence of viral RNA. As expected, in hamsters infected with IAV, no viral RNA could be detected from the surveyed neural tissue that aligned to the IAV genome (Figure S4A). In contrast, within the SARS-CoV-2-infected hamster cohort, viral reads were readily detectable in the nervous system in a subset of animals, consistent with the findings of others (de Melo *et al*., 2021). Of note, in one hamster, SARS-CoV-2 reads were detectable in all surveyed regions of the nervous system (Figure 4B). Mapping of these reads to the SARS-CoV-2 genome revealed that most reads aligned to the nucleocapsid (N) transcript, potentially implicating the deposition of circulating subgenomic RNA from a peripheral infection which is dominated by N (Figure 4B)(Alexandersen et al., 2020).

**Figure 4.**
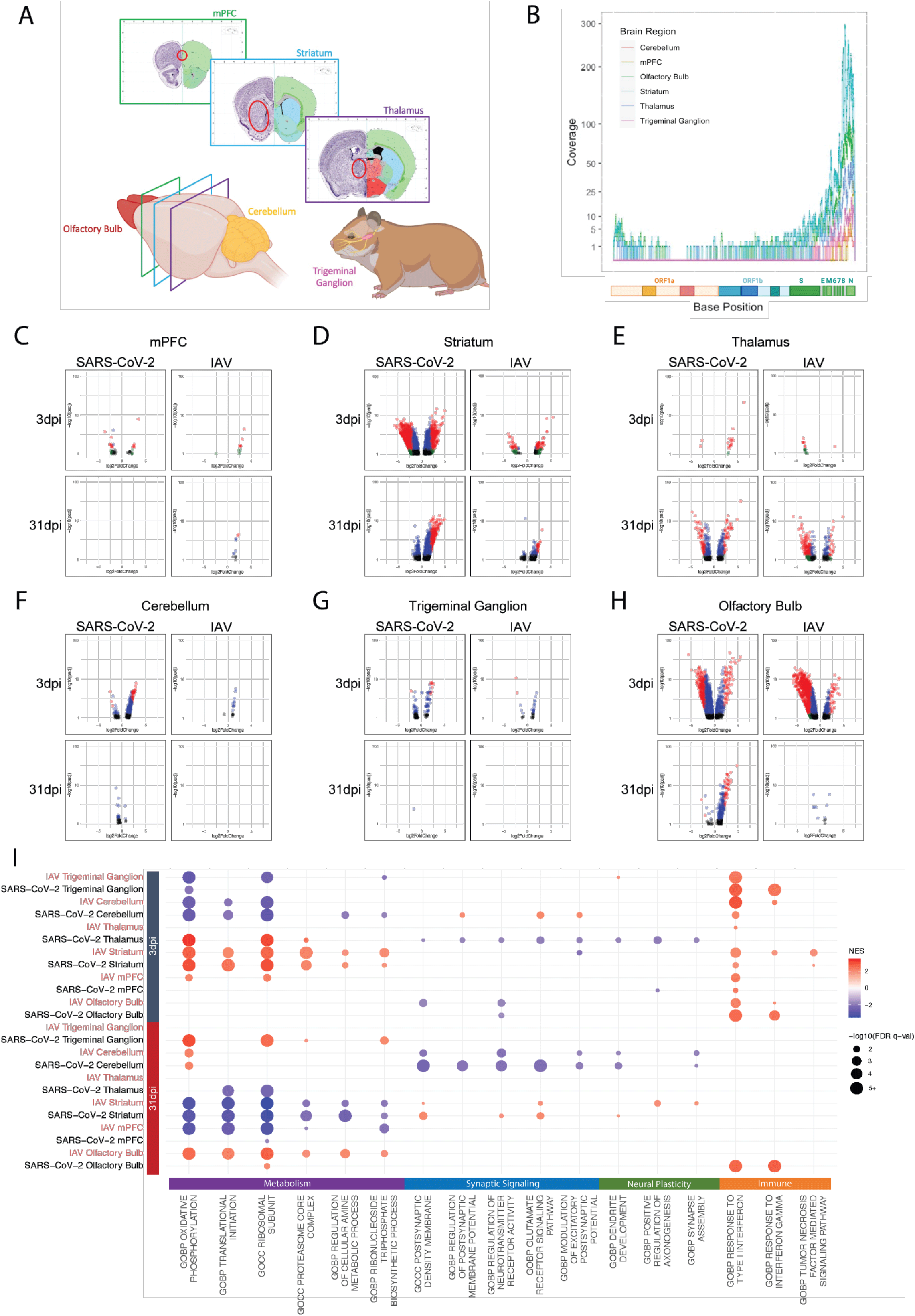
SARS-CoV-2 induces a unique neural transcriptional profile compared to IAV. **(A)** Schematic of brain regions characterized in response to infection. mPFC, striatum, thalamus, cerebellum, trigeminal ganglion, and olfactory bulb were bilaterally harvested for RNA-sequencing analysis. **(B)** Alignment of RNA-Seq data to the SARS-CoV-2 genome. Coverage of raw reads over the length of the genome are displayed as a histogram from each brain region noted. **(C-H)** Differential expression analysis was conducted for IAV- or SARS-CoV-2-infected hamsters compared to mock-infected hamsters at 3dpi and 31dpi via DESeq2; differentially expressed genes with a p-adjusted value of less than 0.1 are plotted (black: p-adj > 0.05, log2 fold-change < 2; blue: p-adj < 0.05, log2 fold-change < 2; green: p-adj > 0.05, log2 fold-change > 2; red: p-adj < 0.05, log2 fold-change > 2). **(I)** Differential expression data analyzed via GSEA using curated gene ontology and human phenotype ontology gene sets; significant enrichments for metabolic-, synaptic signaling-, neural plasticity-, and immune-related ontologies were displayed on the dot plot. Coloration designates normalized enrichment score (NES), with positive enrichment scores colored red and negative enrichment scores colored blue. Dot size is scaled to -log10(FDR q-val) of enrichment; only enrichments with an FDR q-val of less than 0.05 were plotted.

To further characterize the appearance of SARS-CoV-2 genetic material in the brains of infected hamsters, a time course was conducted in which geographically distinct regions of the brain were sampled on days 1, 4, 7, and 14 post-infection in both SARS-CoV-2- and mock-infected hamsters. Olfactory bulb, striatum, and cerebellum were chosen for their respective positioning in the anterior, middle, and posterior sections of the cranial cavity. These regions were assessed for SARS-CoV-2 sgN transcripts via qPCR and compared to lung, the primary site of infection (Figure S4B-E). Mirroring previous data (Figure 1A-B, S1A-B), SARS-CoV-2 sgN detection in the lungs was highest on days 1 and 4. By 7dpi, sgN detection significantly diminished, with only negligible levels detectable at 14dpi (Figure S4B). In the olfactory bulb, a low level of sgN at 1dpi increased to a more prominent level at 4dpi in two of three hamsters before dissipating over the next seven days (Figure S4C). Interestingly, striatum and cerebellum demonstrated different patterns of sgN positivity compared to both lungs and olfactory bulbs. At 1dpi, SARS-CoV-2-infected hamsters demonstrated sgN positivity in one out of three tested striatum sections and in all of the cerebellum samples (Figure S4D-E). Beyond this early time point, however, no cerebellum or striatum sections demonstrated sgN signal that rose above background.

To assess whether sgN positivity was associated with triggering of an innate immune response, transcripts for *Isg15*, a canonical IFN-I stimulated gene, were assessed in sampled regions via qPCR (Figures S4F-I). In general, *Isg15* signal correlated with sgN positivity; in lungs, for instance, *Isg15* signal was elevated on days 1, 4, and 7 post-infection, with its peak at 4dpi (Figure S4F). Striatum and cerebellum likewise show induction of *Isg15* signal at 1dpi in infected tissues, after which *Isg15* expression returns to levels similar to mock-infected tissues; this *Isg15* spike at 1dpi mirrors the positive sgN signal in these tissues (Figures S4D-E and S4G-H). The olfactory bulbs show similar *Isg15* levels following sgN signal on days 1, 4, and 7 post-infection; however, at 14dpi, the olfactory bulb intriguingly shows a newly elevated *Isg15* signal in the absence of any sgN positivity (Figures S4C and S4I).

To better understand the functional impacts that systemic SARS-CoV-2 and IAV challenge have on the nervous system, differential expression analyses of host transcripts were subsequently conducted across all sequenced neural areas from 3 and 31dpi following challenge with either SARS-CoV-2 or IAV and were compared to mock infection (Figure 4C-H). Several brain areas showed significant region-specific transcriptional alterations induced by viral infection at 3dpi. Intriguingly, this was evident for both SARS-CoV-2 and IAV (Figure 4C-H). Direct comparison of SARS-CoV-2 and IAV by differential expression analyses revealed that most surveyed regions induced a very similar transcriptional profile between the two viruses at 3dpi (Figure S4J-K). These findings are most prominent in the striatum, where comparison of SARS-CoV-2 vs. mock conditions revealed more than 3500 DEGs in contrast to the comparison of SARS-CoV-2 vs. IAV, which demonstrate no significant transcriptional differences (Figure 4D and S4J). In contrast, these analyses also revealed significantly different transcriptional profiles in many of the surveyed regions in response to SARS-CoV-2 and IAV infection at 3dpi. These differential responses to the two viral challenges are most prominent in the thalamus, cerebellum, and trigeminal ganglion (Figure 4E-G and S4J-K).

Additionally, this differential expression analysis demonstrated that transcriptional programs induced in neuronal tissue during viral challenge persisted for at least one-month post-infection. Indeed, all surveyed regions showed DEGs in response to at least one of the viruses at 31dpi, albeit strikingly few changes observed in the trigeminal ganglion (Figure 4C-H). These transcriptional signatures were comparable between SARS-CoV-2 and IAV in the striatum, mPFC, and cerebellum (Figure S4J-K). However, similar to the acute response at 3dpi, we again observe specific neuronal regions in which a unique transcriptional signature persists in response to SARS-CoV-2. These virus-specific signatures can be found in the thalamus and olfactory bulb (Figure 4E, 4H, S4J-K).

To better understand the significance of the transcriptional changes taking place during infection, we again performed an unbiased GSEA (Figure 3I). While enrichment of ontologies varied substantially between tissue, timepoint, and viral species, most notable enrichments related to four general categories: metabolism, synaptic signaling, neuronal plasticity, and immune response (Figure 4I). At 3dpi, widespread metabolic modulation was observed within the surveyed neural tissues in response to both SARS-CoV-2 and IAV (Figure 4I, top left quadrant). One example from data collected from the cerebellum and the trigeminal ganglion indicates strong negative enrichment of oxidative phosphorylation. Conversely, the striatum and the thalamus demonstrated inverse trends in response to both SARS-CoV-2 and IAV infections, showing an increase in oxidative phosphorylation amongst other metabolic ontologies. Interestingly, these trends are reversed at 31dpi with cerebellum and trigeminal ganglion showing an increase in oxidative phosphorylation in contrast to striatum and mPFC where these signatures become negatively enriched (Figure 4I). These data suggest a dynamic process of metabolic changes that occur throughout the central nervous system in response to viral challenge.

To better assess the functional impact of viral insult or associated metabolic modulation on neuronal tissue, ontologies relating to synaptic signaling and neural plasticity were further examined. Changes in synaptic signaling showed distinctive responses to SARS-CoV-2 in the thalamus and cerebellum at 3dpi but for all other regions, this response was either not significant or unique when comparing virus challenge models (Figure 4I). Similarly, when examining genes associated with synaptic plasticity, we observe down regulation of these processes predominantly in response to SARS-CoV-2 and primarily only during the acute phase of infection, although unique changes in response to IAV can also be visualized. Lastly, we examined gene ontologies encompassing aspects of the immune response following either SARS-COV-2 or IAV infection. Like the metabolic signatures, we observe similar responses between SARS-CoV-2 and IAV during acute infection. In line with earlier qPCR data from early infection time points (Figures S4F-I), the host response to SARS-CoV-2 or IAV infection results in IFN-I signatures across all neuronal tissues examined with the exception of thalamus (Figure 4I). Despite clearance of virus in both model systems and a broad resetting of neuronal and systemic inflammatory programs across surveyed tissues, we do observe persistent IFN-I and -II signatures selectively in the olfactory bulb following SARS-CoV-2 infection (Figure 4I, bottom right quadrant).

### Characterization of a persistent immune response in the olfactory bulb and epithelium in response to SARS-CoV-2

Given the unique prolonged nature of the proinflammatory response in the olfactory bulb to SARS-CoV-2, we next examined specific genes driving this transcriptional program (Figure 5A-J). Comparing genes implicated in the IFN-I response induced by either IAV or SARS-CoV-2 at 3 and 31dpi highlighted the unique persistence of this transcriptional signature in response to SARS-CoV-2 (Figure 5A). These data demonstrated prolonged elevation of canonical ISGs such as *Isg15*, *Mx2*, and *Irf7* which could be independently corroborated by qRT-PCR (Figure 5B-D). To further confirm these findings, we performed immunostaining for MX1 on sections taken from the olfactory bulbs of hamsters either mock treated or infected with SARS-CoV-2 or IAV at 3 and 31dpi (Figure S5A). These data corroborated our transcriptome findings at the protein level and demonstrated elevated MX1 at both 3 and 31dpi in response to SARS-CoV-2, with immunolabeling remaining in the periphery of the olfactory bulbs (Figure S5A). In addition to ISGs, SARS-CoV-2 infection was also found to induce prolonged chemokine induction as denoted by *Cxcl10* and *Ccl5* amongst others (Figure 5E-F and S5B). Interestingly, the chemokine response observed appears more sustained—or even enhanced at 31dpi—compared to the IFN-I signature.

**Figure 5.**
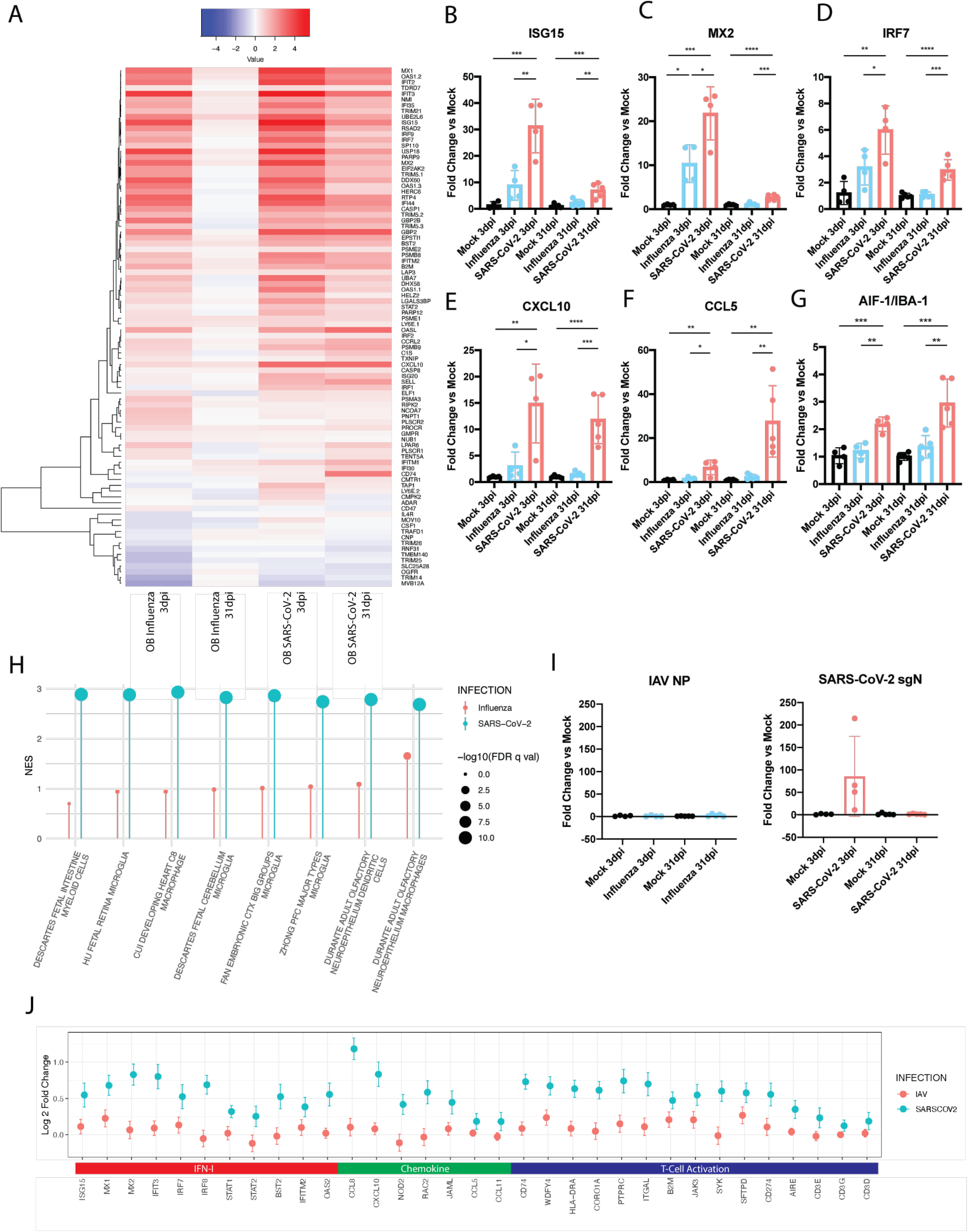
Characterization of persistent inflammation in the olfactory bulb and epithelium in response to SARS-CoV-2. **(A)** Differential expression analysis was conducted on RNA-sequencing of olfactory bulbs of 3dpi and 31dpi IAV- and SARS-CoV-2-infected hamsters compared to mock-infected hamsters. Log2 fold-change of IFN-I response genes are displayed in heatmap. **(B-G)** Expression of key IFN-I response associated genes (**B**) ISG15, (**C**) MX2, and (**D**) IRF7; chemokines (**E**) CXCL10, and (**F**) CCL5; and (**G**) microglial marker AIF-1 (also known as IBA1) were quantified by qRT-PCR. Error bars display standard deviation, and significance was determined by independent tests on samples from 3dpi and 31dpi for each gene using one-way ANOVA with Tukey’s multiple comparisons test (*p<0.05, **p<0.01, ***p<0.001, ****p<0.0001). **(H)** Differential expression data from 31dpi olfactory bulb RNA-sequencing was analyzed by GSEA analysis using MSigDB C8 gene set collection, which contains gene sets derived from single cell RNA-sequencing datasets that define specific cellular identity signatures. Top positive enrichments are plotted by their normalized enrichment score (NES) (magnitude of line) and significance (-log10(FDR q-val)) (dot size). **(I)** Levels of IAV or SARS-CoV-2 RNA in the olfactory bulbs as measured by RT-qPCR with primers for IAV nucleoprotein (IAV NP) or SARS-CoV-2 subgenomic nucleocapsid protein (SARS-CoV-2 sgN), respectively. Error bars display standard deviation. **(J)** Olfactory epithelium from mock-, IAV-, or SARS-CoV-2-infected hamsters were harvested and transcriptionally profiled via RNA-sequencing. Differential expression analysis was conducted on infected groups compared to mock. Log2 Fold Change for expression of individual genes relevant to ontologies concerning IFN-I, chemokine, and T cell activation are presented in the graph for both IAV and SARS-CoV-2 compared to mock; error bars denote standard error.

We next sought to determine the composition of immune cells participating in the prolonged inflammatory response. To this end, we analyzed transcriptomic data from olfactory bulbs at 31dpi to identify enriched gene sets that enable deconvolution to identify specific cell types present. These analyses showed profound enrichment for microglial and myeloid lineage gene sets, specifically in the SARS-CoV-2-infected hamsters at this time point (Figure 5H). This analysis was further supported by a directed GSEA which demonstrated a significant positive enrichment in markers for microglial activation (Figure S5C). To better assess how CNS-specific cell types were changing in response to SARS-CoV-2 infection at this time point, gene sets were created for neuronal and glial cell populations using known cell-type markers identified previously (Zhang et al., 2014). These efforts confirmed significant positive enrichment of microglial-specific transcriptomic signatures in olfactory bulbs from SARS-CoV-2-infected hamsters at 31dpi (Figure S5D). In contrast to immune cells, gene sets identifying neuronal populations demonstrated significant negative enrichment at this same time point, indicating a potential loss of, or altered activity within, the neurons of SARS-CoV-2-infected hamsters as compared to mock-infected controls.

To further corroborate the transcriptional signatures of the olfactory bulbs in response to either SARS-CoV-2 or IAV, we performed independent qPCR validation and/or immunohistochemistry on genes for which commercial antibodies for the hamster were available. These efforts illustrated a significant recruitment of microglial/macrophage populations to the olfactory bulbs, as measured by *Aif-1* transcripts, uniquely in response to SARS-CoV-2 (Figure 5G). We next aimed to assess microglial activation at the histological level. Immunostaining for IBA-1, the protein encoded by *Aif-1*, demonstrated increased microglial positivity around the periphery of the olfactory bulb. This finding was most pronounced in olfactory bulbs from SARS-CoV-2-infected hamsters at 3dpi but could still be seen at 31dpi (Figure S5E-F). Microglia could be visualized clustering around the periphery of the olfactory bulbs in both SARS-CoV-2- an IAV-infected hamsters (Figure S5F-G). While the histological analyses revealed comparable IBA-1 staining between SARS-CoV-2 and IAV, it should be noted that microglial activation results in significant transcriptional changes following activation which were uniquely prominent in response to SARS-CoV-2 (He et al., 2021).

While GSEA data primarily implicated microglial and myeloid lineages in the inflammatory phenotype, a milder positive enrichment of T cell signatures was also noted. To assess the degree of T cell infiltration involved in the hamster response, olfactory bulbs from 31dpi were immunolabeled for CD3 (Figure S6A). Staining was noticeably sparse, with minimal numbers of cells (∼20-50 cells/bulb) labeled positive in mock, IAV, and SARS-CoV-2 olfactory bulb cross-sections.

To determine if sustained IFN-I or chemokine expression was the product of a chronic infection, we next performed qRT-PCR on the olfactory bulbs (Figure 5I). These data clearly demonstrate that at 31dpi, neither IAV nor SARS-CoV-2 transcripts could be detected, although SARS-CoV-2 subgenomic Nucleocapsid (N) levels were evident at 3dpi. qRT-PCR data was further validated by RNA *in situ* hybridization which confirmed that SARS-CoV-2 Spike (S) staining was only observed at 3dpi in the glomerular region and was undetectable at 31dpi (Figures 5I and S6B).

To further assess whether inflammation of the olfactory bulbs was associated with cellular apoptosis in this region, we performed a TUNEL stain on mock-, SARS-CoV-2-, or IAV-infected hamsters at these times. Quantification of the total number of TUNEL-positive nuclei in the olfactory bulbs revealed no significant differences between the infection groups at either time point, indicating that neither the acute infection, nor the uniquely prolonged inflammatory patterns in SARS-CoV-2 olfactory bulbs, were associated with local apoptosis (Figure S6C).

To explore whether this proinflammatory signal was present in additional anatomical regions linked to the olfactory bulb, olfactory epithelium was harvested from hamsters infected with SARS-CoV-2, IAV, or PBS (mock) at 30dpi as this tissue has been demonstrated to harbor infectious virus (Hoagland *et al*., 2021; Horiuchi et al., 2021). Ontological analyses of mRNA-Seq data demonstrated that, similar to the olfactory bulbs, the olfactory epithelium of SARS-CoV-2-infected hamsters uniquely showed upregulated signatures for IFN-I and IFN-II (Figure 5J and S6D). These signatures were driven by expression of canonical ISGs such as *Isg15, Mx1, Mx2, Ifit3, Irf7, Oas2* and *Bst2*. Intriguingly, however, ontological analysis also highlighted several transcriptomic signatures implicating T cell recruitment, activation, differentiation, and immune response in the olfactory epithelium. Chemotactic recruitment signatures were driven by increases in expression of genes such as *Ccl7, Cxcl10, Ccl5,* and *Ccl11* as well as other cellular migration factor genes, such as *Jaml* and *Rac2* (Figures 5J and S6D). T cell activation ontologies, on the other hand, were driven by upregulated expression of antigen presentation markers, such as *Hla-dra, Wdfy4,* and *B2m* concurrently with upregulation of T cell-associated genes such as *Jak3, Coro1A, Cd3e, Cd3g,* and *Cd3d* (Figures 5J and S6D). In addition to immune signatures, these analyses highlighted a negative enrichment for genes relating to sensory perception of smell and olfaction capabilities which were present for both SARS-CoV-2- and IAV-infected hamsters (Figure S6D).

To better understand the cellular make up of this immune response, cell type enrichment analyses were again conducted (Figure S6E). Enrichment analyses implicated the presence of unique neuroepithelium lymphocytes and macrophage populations in the olfactory epithelium following SARS-CoV-2 infection.

Importantly, as SARS-CoV-2 has demonstrated sex-dependent biases, we also assessed whether evidence for sustained perturbations in the olfactory bulbs and/or epithelium were present in female hamsters. To this end, a cohort of all female hamsters were infected with SARS-CoV-2 or IAV and analyzed at 24dpi (Figure S6F-G). Consistent with our earlier results performed in male hamsters, we find elevated Isg15 and Ccl5 levels in both tissues.

### Olfactory inflammation is associated with behavioral alteration

Given prior findings that hamsters, similar to humans, can experience anosmia in response to SARS-CoV-2 infection (de Melo et al., 2021), and the fact that injury to the olfactory bulb has been linked to development of neurobehavioral disorders such as depression (Hasegawa-Ishii et al., 2019; Hellweg et al., 2007; Kelly et al., 1997; Kim et al., 2019; Song and Leonard, 2005), we next strove to assess the functional consequences of sustained neuronal perturbations, such as prolonged olfactory bulb and epithelium inflammation in SARS-CoV-2-infected hamsters beyond 4 weeks post-infection.

To this end, we first looked to elucidate how SARS-CoV-2 infection affected olfaction. Hamsters infected with IAV or SARS-CoV-2 were assessed for smell and compared to a cohort of mock-infected animals. Utilizing a food-finding test at 3dpi, 15dpi, and 28dpi, we confirmed the results reported in de Melo et al., showing SARS-CoV-2-infected hamsters took longer to find buried food at 3dpi while showing no significant difference at 15 or 28dpi when compared to mock or IAV which were indistinguishable and were thus plotted together (de Melo *et al*., 2021)(Figure 6A-F and S6H-J). In contrast, when this same experiment was performed with readily visible food, all cohorts, at all time points tested, displayed roughly equivalent times (Figure S6H-J). This result came despite prolonged inflammation in both olfactory bulb and epithelium and coincided with transcriptional signatures indicative of diminished sensory perception of smell for both IAV and SARS-CoV-2 (Figure S6D).

**Figure 6.**
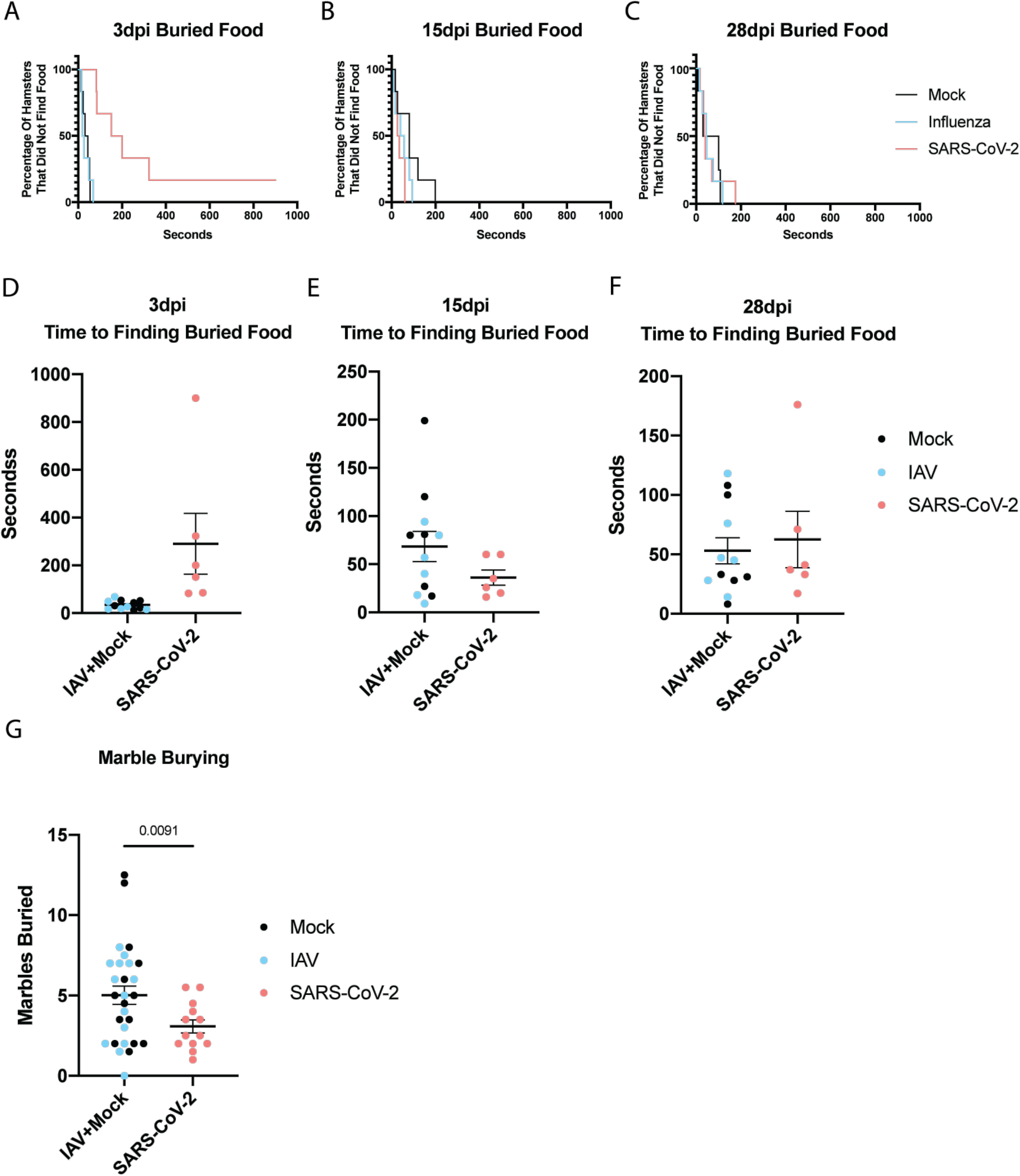
Olfactory inflammation is associated with behavioral alteration. **(A-F)** IAV-, SARS-CoV-2, and mock-infected hamsters were assessed for smell at **(A)** 3dpi, **(B)** 15dpi, and **(C)** 28dpi using a buried food finding test. Kaplan-Meier curves demonstrate time to discovery of food across all time points and infection groups. Throughout the analyses, IAV- and mock-infected hamsters displayed nearly identical phenotypes, so they were grouped together to better highlight changes in SARS-CoV-2 infection group performance at (**D)** 3dpi, **(E)** 15dpi, and **(F)** 28dpi as measured by the time it took hamsters to find the buried food. **(G)** All infection groups were assessed for behavior at 26dpi using the marble burying assay, a test classically utilized to measure repetitive, obsessive-compulsive, and anxiety-like behavior in rodents. The number of marbles that were greater than 60% buried were counted and graphed for each hamster.

To determine whether prolonged olfactory bulb inflammation was correlated with altered metrics on assays that assess affective behaviors, mock-, IAV-, and SARS-CoV-2-treated hamsters were subjected to a marble burying assay (Figure 6G-I). When hamsters were subjected to the this assay, an established metric for assessing rodent repetitive and anxiety-like behaviors (Yanai and Endo, 2021), SARS-CoV-2-infected animals demonstrated a significant reduction in burying activity compared to both mock and IAV groups, which performed comparably and were thus grouped together (Figure 6G). While several studies consider increased burying as a sign of elevated compulsiveness or anxiety-like behaviors, other studies have found that anxiogenic or compulsive behavior-inducing pharmacological substances can actually cause decreases in marble burying behavior that are well-below baseline (Jimenez-Gomez et al., 2011). In light of bi-directionally affected marble burying behavior suggesting altered affective states, this data suggests that SARS-CoV-2 may induce mild, yet significant behavioral changes, and result in a mild vulnerability to environmental stressors.

### SARS-CoV-2 infection is associated with sustained inflammatory transcriptional programs in human olfactory bulb and olfactory epithelium

Finally, to ascertain whether our data could be extended to aspects of the human disease, we performed RNA sequencing on post-mortem olfactory bulb and olfactory epithelium tissue from human donors that had recovered from a medically documented history of COVID-19 infection that had occurred at least one month prior to death (Figure 7A-F). Donors were screened to include only those who had died of causes unrelated to COVID-19 disease. Tissues from healthy donors without history of COVID-19 infection were also collected as controls.

**Figure 7.**
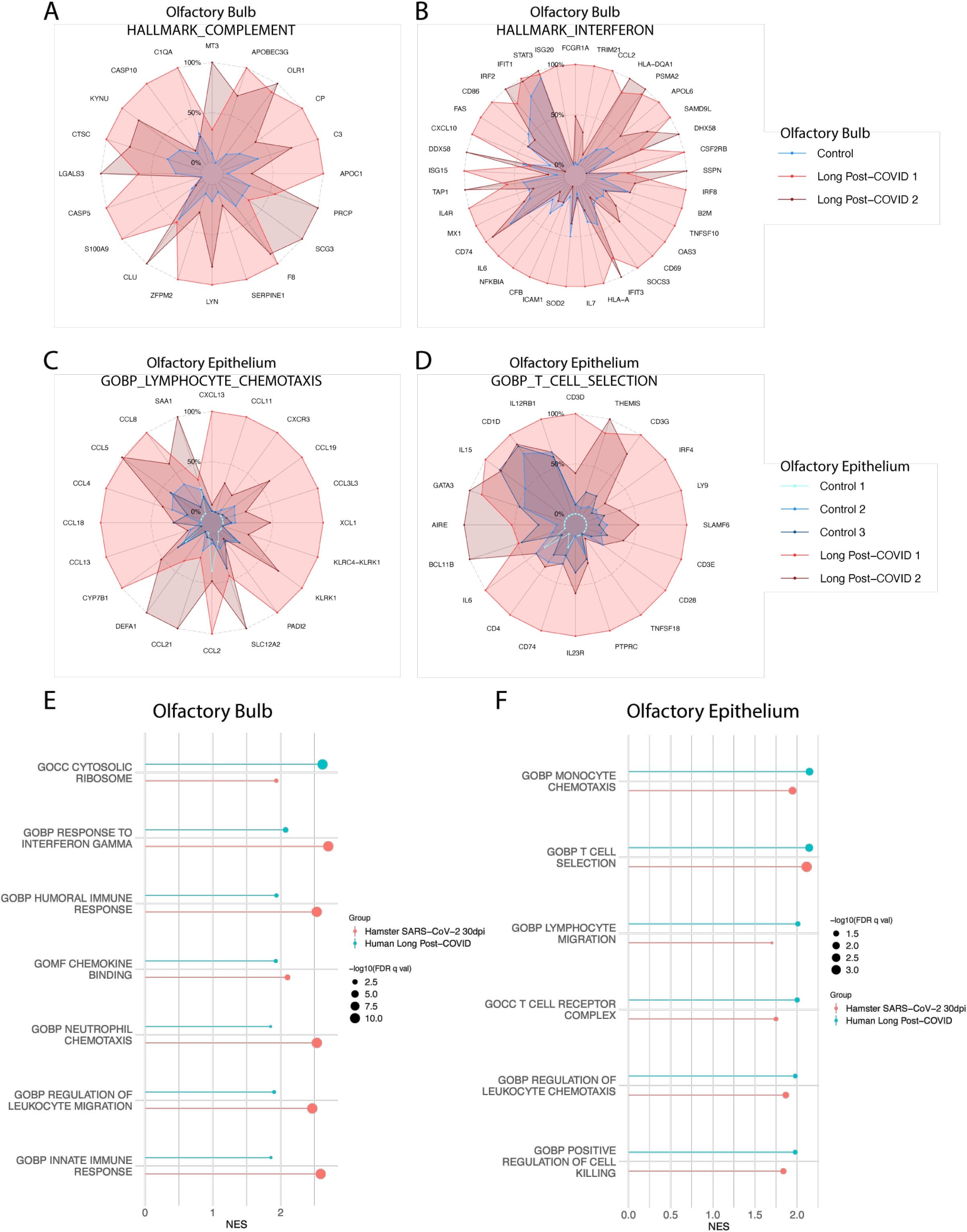
SARS-CoV-2 infection is associated with sustained inflammatory transcriptional programs in human olfactory bulb and olfactory epithelium. **(A-B)** Radar plots derived from olfactory bulb tissues collected at autopsy from healthy control donors (Control) as well as donors that had previously recovered from clinically documented COVID-19 (Long Post-COVID). Donors were screened to only include those where COVID-19 positivity was documented >1 month prior to autopsy. Tissues were RNA-sequenced, and Long Post-COVID tissues were compared to control tissues via differential expression analysis. GSEA using the Hallmark Gene sets was utilized to characterize transcriptomic programs. Transcripts per million reads (TPM) counts for individual genes making up these responses were plotted onto radar plots. Gene expression is normalized to the highest expressing sample for each individual gene, with expression levels shown as the percentage of TPM value of this sample (which is shown as 100% of its own value). **(C-D)** Analyses as described in (A) were used to characterize the transcriptional response of olfactory epithelium tissues harvested from Long-Post COVID and control donors. **(E-F)** GSEA enrichment analyses from **(E)** olfactory bulb and **(F)** olfactory epithelium human tissues were plotted by their normalized enrichment scores (NES) (magnitude of line) and significance (-log10(FDR q-value)) (size of dot). GSEA enrichments of these same gene sets from analogous tissue analysis in hamsters (SARS-CoV-2-infected vs mock-infected olfactory bulb and epithelium tissues at 31dpi) was plotted beside matching human enrichment data.

For olfactory bulb tissues, two COVID-19 recovered (long post-COVID) donors and one uninfected control donor were able to be sequenced. Differential expression and gene enrichment analyses revealed the presence of proinflammatory transcriptional programs in the olfactory bulbs of the recovered donors compared to control tissues (Figure 7A-B). In agreement with that observed in hamsters, gene sets for complement (Figure 7A) and interferon (Figure 7B) demonstrated significant enrichment. Complement gene set enrichment was driven by upregulation of direct complement cascade genes such as *C3, F8,* and *C1QA* as well as complement regulatory proteins such as *S100A9, SERPINE1, and CLU* (Figure 7A). Interferon ontologies were driven by shared upregulation of ISGs such as *ISG15, OAS3, ISG20, CXCL10, MX1, IFIT3, IFIT1,* and *IRF2*, as well as other immune-related genes such as those involved in antigen presentation (*B2M, HLA-DQA1, CD74*) and cytokine signaling (*IL7, IL6*, *IL4R*) (Figure 7B). Of note, one long post-COVID donor (Long Post-COVID 1) showed a higher level of inflammatory gene expression than the other, possibly reflecting their medical history as this individual had COVID-19 within four months of death as opposed to the companion sample (∼six months).

For olfactory epithelium tissues, two long post-COVID donors and three uninfected control donors were able to be successfully sequenced. Differential expression and gene enrichment analyses showed that, similar to findings in olfactory bulb tissues, long post-COVID donor tissue displayed enriched proinflammatory transcriptional profiles (Figure 7C-D). However, while tissue from olfactory bulbs were dominated by interferon-mediated transcriptional programs, olfactory epithelium tissues displayed a higher enrichment for gene sets detailing chemotactic and T cell-specific activities. These programs were respectively driven by upregulation of a variety of chemotactic (i.e. *CCL5, CCL8, CCL19, CXCR3*) and T cell-associated genes (i.e. *CD3D, CD3G, CD3E, GATA3, CD4, LY9*) (Figure 7C-D). Once again, one long post-COVID donor (Long Post-COVID 1) appeared to have a higher level of chemotactic and T cell markers than the other long post-COVID sample; likely reflecting the time between COVID-19 and death.

Comparison of the olfactory bulbs and olfactory epithelium from 31dpi derived from SARS-CoV-2-infected hamsters and long post-COVID humans demonstrated a strong degree of concordance between the respective transcriptional programs. In the olfactory bulb, both hamster and human SARS-CoV-2-recovered tissues show enhanced induction of IFN-II, leukocyte chemotaxis, and immune response pathways (Figure 7E). The two organs further show coordinated metabolic programs, with both hamster and human post-SARS-CoV-2 infection tissues demonstrating upregulation of ribosomal production (Figure 7E). Moreover, in the olfactory epithelium, both human and hamster tissues demonstrate enrichment of chemotaxis and T cell functional pathways after SARS-CoV-2 clearance. The strong correlation generated when comparing transcriptional responses between hamsters and humans that have recovered from SARS-CoV-2 would suggest that the host response results in prolonged olfactory inflammation in both infections.

## DISCUSSION

Together, these data demonstrate that SARS-CoV-2 and IAV infections produce a wide range of longitudinal systemic impacts that include both shared and unique characteristics between the two viruses. In peripheral tissues, such as lung, heart, and kidney, SARS-CoV-2 and IAV seem to induce similar transcriptomic and histological changes both during active infection and following viral clearance. However, levels of scarring from infection were more severe following SARS-CoV-2 which induced a higher degree of kidney tubular atrophy and lambertosis compared to IAV. These differences likely reflect the unique biologies of the viruses. SARS-CoV-2 generates significantly more double stranded RNA (dsRNA) during its life cycle as a result of sgRNA production (Nilsson-Payant *et al*., 2021; Perlman et al., 1986). Given the immunogenicity and stability of dsRNA, it seems reasonable to postulate that comparable levels of replication would result in a more robust immune response to SARS-CoV-2 as compared to IAV. Perhaps it is the unique magnitude of the host response to SARS-CoV-2 that induces the observed pathological abnormalities which would result in reduction of functional capacity in affected regions, as supported by previously reported data (Allen, 2010; Eddy, 2005; Fukuoka et al., 2005; Schelling, 2016; Wright *et al*., 2020; Yamashita et al., 2020). Similar to peripheral tissues, the nervous system showed a mix of shared and unique responses to SARS-CoV-2 and IAV infection. During acute infection, both viruses induced central nervous system (CNS)-wide IFN-I responses as well as region-specific transcriptional alterations that in some cases persisted beyond one month following infection. The most prominent of these alterations took place in the striatum, where *both* IAV and SARS-CoV-2 induced similar changes associated with metabolic and functional shifts. Pre-clinical and clinical literature have correlated this type of activity within striatal subregions with chronic or traumatic stress (Magalhaes et al., 2019; Rangaprakash et al., 2017), affective disorders (Oathes et al., 2015; Torres-Sanchez et al., 2017), and chronic pain states (Serafini et al., 2020). These changes could partially underlie the increased clinical risk of neurological and neuropsychiatric disorder onset associated with both IAV (Bornand et al., 2016) and SARS-CoV-2 (Huang et al., 2021; Taquet et al., 2021).

In contrast to the striatum, the thalamus displays a differentially regulated response to SARS-CoV-2 and IAV infection. At 3dpi, SARS-CoV-2 infection induces a hypoexcitable state in the thalamus, while IAV pushes the region towards hyperexcitability. Clinical literature has identified cognitive deficits (Hosp et al., 2021) associated with acute, severe COVID-19, including confusion and dysexecutive syndrome (Beaud et al., 2021). While these are generally attributed to abnormalities in the nuclei of the frontal lobe, we found little evidence of transcriptional dysregulation in the mPFC, a key executive region. Our sequencing and ontological analyses suggests instead that thalamic dysregulation may contribute to cognitive disruption, potentially in the form of altered intra-thalamic function or functional connectivity with key brain regions that drive emotion, motivation, cognition, sleep, pain, wakefulness, and motor activity. Altered thalamic function and structure has been previously associated with cognitive deficits in conditions such as multiple sclerosis (Schoonheim et al., 2015), traumatic brain injury (Grossman et al., 2012), and Alzheimer’s disease (Wang et al., 2012). Thalamic dysfunction may also underlie neurological conditions that have been observed in long COVID patients including chronic pain, headache, myalgias, seizures, sleep, and affective disorders (Feng et al., 2017; Greicius et al., 2007; Gustin et al., 2014; Hodaie et al., 2002; Iadarola et al., 1995; Li et al., 2019; Noseda et al., 2017). Furthermore, transcriptomic changes strongly associated with dendrite development were also seen in this region at both early and late time points after SARS-CoV-2-infection but not in response to IAV-infection. Intriguingly, dysregulation of the key genes driving this enrichment in the SARS-CoV-2 thalamus (*Nrp1, App, Crtc1, Ctnnd2, Camk2a, Kalrn, Bmp7, Ppp1r9b, Mecp2, Cux1, Dlg4, Apoe, Ephb2, Map2k7, Ephb1)* are associated with cognitive impairments and affective disorders such as major depressive disorder when analyzed together using Enrichr’s DisGeNET function (Kuleshov et al., 2016). Thus, regional transcriptional changes in thalamic nuclei may facilitate the development of neuropsychiatric disorders in patients recovering from SARS-CoV-2 infection.

By far, the most unique response to SARS-CoV-2 took place in the olfactory bulb. At 31dpi, the olfactory bulb of IAV-infected hamsters returned to a baseline transcriptional state while the same region of SARS-CoV-2-infected hamsters appeared to be in the midst of an ongoing infection characterized by microglial activation and a robust IFN-I and chemokine response. This was especially surprising given that we were unable to detect presence of viral RNA in either the olfactory bulb or lungs at this time point and, demonstrated previously, we know that hamsters generate a strong anti-S antibody response as early as seven days post infection (Hoagland *et al*., 2021; Horiuchi *et al*., 2021). Immunohistochemistry of MX1 corroborated our transcriptional findings and showed localization of this persistent IFN-I response to the glomerular regions of the olfactory bulbs. Given the olfactory bulbs were positive for SARS-CoV-2 early in infection, these data may suggest either an undetectable level of replication is persisting in an adjacent region or that left-over debris is responsible for the continued inflammatory profile. Our inability to isolate infectious virus from hamsters from any tissue after seven days post infection would provide support for the latter hypothesis, although others have reported the ability to detect replication-competent virus from comparable tissues (de Melo *et al*., 2021; Hoagland *et al*., 2021; Horiuchi *et al*., 2021).

Whatever the cause, chronic inflammation within the olfactory bulbs could impact sensory, emotional, and cognitive processes. The olfactory bulbs are functionally connected to—and can thus influence activity of—the limbic system, which controls appetitive, sensory, emotional, and cognitive responses. Indeed, prior preclinical studies link olfactory bulb damage with depressive phenotypes that can be reversed with antidepressant treatment (Guarnieri et al., 2020; Hellweg et al., 2007; Kelly et al., 1997; Song and Leonard, 2005). These data suggest that chronic nasal and olfactory bulb inflammation may drive neurodegeneration and structural changes consistent with long COVID symptoms (Hasegawa-Ishii et al., 2017; 2019). This is further supported by recently reported clinical evidence that shows that patients that have recovered from even mild COVID-19 demonstrate loss of grey matter in limbic cortical areas functionally linked to the olfactory system (Douaud et al., 2021). Taken together, our peripheral organ and central nervous system findings identify transcriptional and histologic signatures caused by SARS-CoV-2 infection that may induce a variety of somatosensory, affective, and cognitive impairments that persist well past the time of original infection. Given the systemic scope of these findings, we hypothesize that they elucidate a molecular basis of much of the heterogenous symptomology that makes up long COVID.

## ACKNOWLEDGEMENTS

This work was funded by generous support from the Zegar Family Foundation to the B.T. and funding from NINDS (NS111251, NSO86444, NSO86444S1 to V.Z. and R.A.S. In addition to funding support, we are indebted to the patients, their families, and healthcare workers that have been fighting the COVID-19 pandemic. In addition, we would like to thank Steven Salvatore, MD, for sharing his clinical expertise and for his work reviewing our kidney histology. We would like to further thank Francis Avila, Virginia Gillespie, DVM, Ying Dai, and the rest of the staff at the Mount Sinai Center for Comparative Medicine and Surgery and the Mount Sinai Biorepository and Pathology Core for their technical assistance in tissue preparation for histology. We would finally like to thank Alfred D. Doyle, PhD, for his technical guidance. Sequencing of some samples were performed at the New York Genome Center (NYGC) as part of the COVID-19 Genomic Research Network (CGRN) with funds generously provided by NYGC donors.

## AUTHOR CONTRIBUTIONS

Project was conceptualized by J.J.F., R.A.S., V.Z., and B.T. Data was curated by J.J.F., M.Z., S.L., and B.T. Investigation was conducted by J.J.F., R.A.S., K.D.P., M.Z., K.O., M.P., I.G., J.Z., S.H., D.A.H., R.M., A.R., A.K., Human OE/OB tissue was obtained from J.B.O, P.D.C., and J.E.G. Formal analysis was conducted by J.J.F. and A.C.B. Project administration was overseen by J.J.F., R.A.S., R.S, V.Z., and B.T. Writing—original draft was done by J.J.F., R.A.S., and B.T. Additional Writing— review & editing was performed by V.Z. Funding acquisition was done by S.L., V.Z. and B.T.

## SUPPLEMENTAL INFORMATION LEGENDS

**Figure S1 SARS-CoV-2 and IAV infections in hamsters induce transcriptional and histological changes that mirror human infection pathology**

**(A-B)** Lungs of an independent longitudinal cohort of hamsters infected with (**A**) IAV or (**B**) SARS-CoV-2 were measured for viral load via RT-qPCR with primers for IAV nucleoprotein (IAV NP) or SARS-CoV-2 subgenomic nucleocapsid protein (SARS-CoV-2 sgN), respectively.

**(C-D)** H&E staining was conducted on (**C**) kidneys and (**D**) hearts of SARS-CoV-2-, IAV-, and mock-infected hamsters at 3dpi. Histological analysis of hearts confirmed by board-certified pathologist revealed leukocytic infiltration (green stars).

**(E-F)** Volcano plots depicting differential expression analysis conducted on RNA-sequencing data derived from lungs of IAV- or SARS-CoV-2-infected hamsters compared to mock-infected hamsters at 3dpi via DESeq2; differentially expressed genes with a p-adjusted value of less than 0.1 are plotted (black: p-adj > 0.05, log2 fold-change < 2; blue: p-adj < 0.05, log2 fold-change < 2; green: p-adj > 0.05, log2 fold-change > 2; red: p-adj < 0.05, log2 fold-change > 2).

**(G)** Lollipop charts denoting differential expression data analyzed via GSEA using the Hallmark gene sets. Charts display normalized enrichment score (NES) and a dot size scaled relative to -log10(FDR q-val) of the enrichment for top ten most positively enriched gene sets and top three most negatively enriched gene sets in this analysis.

**Figure S2 SARS-CoV-2 and IAV induce lasting transcriptional signatures in peripheral organs that are detectable at 31dpi**

**(A-C)** Volcano plot denoting RNA-sequencing data conducted on **(A)** lungs, **(B)** hearts, and **(C)** kidneys tissue of IAV-, SARS-CoV-2-, and mock-treated hamsters at 31dpi. Differential expression analysis was computed using DESeq2; differentially expressed genes with a p-adjusted value of less than 0.1 are plotted (black: p-adj > 0.05, log2 fold-change < 2; blue: p-adj < 0.05, log2 fold-change < 2; green: p-adj > 0.05, log2 fold-change > 2; red: p-adj < 0.05, log2 fold-change > 2).

**(D)** RNA-sequencing data for all heart, lung, and kidney samples were hierarchically clustered by maximal distance between read data for each sample.

**(E-G)** GSEA analysis using the MSigDB C5 curated gene ontology set was conducted on 3dpi IAV vs. Mock and 3dpi SARS-CoV-2 vs. Mock differential expression data for **(E)** lungs, **(F)** heart, and **(G)** kidneys. Top significant ontological enrichments in SARS-CoV-2-associated analyses are plotted by their normalized enrichment score (NES) (line magnitude) and significance (-log10(FDR q-value)) (dot size). GSEA enrichment for these same gene sets in IAV vs. Mock differential expression data for the same tissue is plotted side-by-side for comparison.

**(H-J)** GSEA using the Hallmark gene sets was conducted on RNA-sequencing differential expression data comparing human early-infection SARS-CoV-2 lung tissue from post-mortem donors to control human tissues for enriched ontologies listed. Heat maps show log2 Fold Change of individual genes comprising the respective gene sets for both hamster and human infection groups compared to control tissues.

**Figure S3 Histological analyses of SARS-CoV-2 and IAV-infected hamsters at 31dpi**

**(A-C)** Immunohistochemical labeling for **(A)** IBA1, **(B)** MPO, and **(C)** CD3, were used to label macrophage, neutrophil, and T cell populations, respectively, in the lungs of mock-, IAV-, and SARS-CoV-2-infected hamsters at 31dpi. Size of inset scale bars matches length described in column headers.

**(D)** Verhoeff Van Gieson staining was performed on sections derived from the lungs of 31dpi SARS-CoV-2-, IAV-, and mock-infected hamsters.

**(E)** H&E staining was conducted on hearts of SARS-CoV-2-, IAV-, and mock-infected hamsters at 31dpi.

**(F)** Airway sizes in cross-sections of lungs from 31dpi SARS-CoV-2-, IAV-, and mock-infected hamsters were quantified via morphometric image analysis. Error bars display standard error mean, and significance was quantified via one-way ANOVA with Dunn’s Multiple Comparison Test (*p<0.05, **p<0.01, ***p<0.001)

**Figure S4 SARS-CoV-2 and IAV induce both unique and shared transcriptional responses in the nervous system**

**(A)** Olfactory bulbs, mPFC, striatum, thalamus, cerebellum, and trigeminal ganglion were bilaterally sampled for RNA-sequencing analysis from 3dpi IAV-infected hamsters. Sequencing reads were aligned to the IAV A/California/04/2009 genome, and coverage of raw reads over the length of the genome were displayed as a histogram for each brain region from the hamster with the highest number of reads.

**(B-E)** Lung, olfactory bulbs, striatum, and cerebellum were sampled from a longitudinal SARS-CoV-2- or mock-infected hamster cohort at 1, 4, 7, and 14dpi and assessed for SARS-CoV-2 subgenomic nucleocapsid (sgN) protein transcripts via qRT-PCR. Values shown have Ct values for actin (*Actb*), a housekeeping control gene, subtracted from Ct values for sgN for normalization.

**(F-I)** These longitudinal tissues were also assessed for expression of canonical IFN-I gene *Isg15* via qRT-PCR. Values shown display fold change normalized to mock controls.

**(J)** Differential expression analysis was conducted for SARS-CoV-2-infected hamsters compared directly to IAV-infected hamsters at 3dpi and 31dpi via DESeq2 across the sampled brain regions; differentially expressed genes with a p adjusted value of less than 0.1 are plotted (black: p-adj > 0.05, log2 fold-change < 2; blue: p-adj < 0.05, log2 fold-change < 2; green: p-adj > 0.05, log2 fold-change > 2; red: p-adj < 0.05, log2 fold-change > 2).

**(K)** Rank-rank scatter plots were generated for each brain region at 3dpi and 31dpi to compare coordination of gene expression from SARS-CoV-2- and IAV-infected hamsters when compared to mock-infected hamsters.

**Figure S5 SARS-CoV-2 induces a uniquely prolonged chemokine signature detectable at 31dpi in the olfactory bulb**

**(A)** Formalin-fixed paraffin-embedded (FFPE) olfactory bulbs from mock-, IAV-, or SARS-CoV-2-infected hamsters at 3dpi and 31dpi were immuno-labeled for MX1 protein. Zoomed inset displays glomerular area of maximum positivity within each sample.

**(B)** Differential expression analysis was conducted on RNA-sequencing of olfactory bulbs of 3dpi and 31dpi IAV- and SARS-CoV-2-infected hamsters compared to mock-infected hamsters. Log2 fold-change of curated chemokine genes from this analysis are displayed in heatmap.

**(C)** GSEA of the GOBP_MICROGLIAL_CELL_ACTIVATION ontology was conducted on RNA-sequencing differential expression data from olfactory bulbs of 3dpi and 31dpi IAV- and SARS-CoV-2-infected hamsters and displayed as a GSEA enrichment plot.

**(D)** Differential expression data comparing olfactory bulbs of 3 and 31dpi infected hamster cohorts to mock-infected was analyzed for enrichment of cell types via GSEA. Gene sets surveyed in this analysis were created using characterized cell type-specific markers characterized in Zhang et al. (2014). Normalized enrichment score (NES) is plotted. Enrichments achieving significance (FDR q-value < 0.05) are outlined in black.

**(E-F)** Formalin-fixed paraffin-embedded (FFPE) olfactory bulbs from mock-, IAV-, or SARS-CoV-2-infected hamsters at **(E)** 3dpi and **(F)** 31dpi were immuno-labeled for IBA-1 protein. Inset scale bar is representative of the length displayed at the bottom of the given column.

**Figure S6 SARS-CoV-2-infection is associated with olfactory epithelium inflammation and anosmia that resolves over time**

**(A)** Formalin-fixed paraffin-embedded (FFPE) olfactory bulbs from mock-, IAV-, or SARS-CoV-2-infected hamsters at 31dpi were immuno-labeled for CD3 protein. Inset scale bar is representative of the length displayed at the bottom of the given column.

**(B)** In-situ hybridization for SARS-CoV-2 S protein was conducted on FFPE olfactory bulbs from SARS-CoV-2-infected hamsters at 3dpi and 31dpi. S protein transcripts are visible as yellow puncta and DAPI nuclear staining is visible in blue. Zoomed inset displays representative glomerular area of high positivity within each sample.

**(C)** Formalin-fixed paraffin-embedded (FFPE) olfactory bulbs from mock-, IAV-, or SARS-CoV-2-infected hamsters at 31dpi were processed via TUNEL staining to assess for apoptotic cells. The number of positive cells manually quantified in each olfactory bulb section is reported here.

**(D)** Olfactory epithelium from SARS-CoV-2-, IAV-, and mock-infected hamsters was harvested at 31dpi and assessed via RNA-seq. Differential expression analysis was conducted with DESeq2 and analyzed via Gene Set Enrichment Analysis (GSEA) using curated gene ontology sets. Lollipop charts show significance of enrichment (- log10[FDR q-val]) (dot size) and normalized enrichment score (NES) for SARS-CoV-2 and IAV vs. mock.

**(E)** GSEA was also performed on 31dpi olfactory epithelium differential expression data to assess for enrichment of cell-specific transcriptional signatures in SARS-CoV-2- and IAV-infected hamsters compared to mock.

**(F-G)** Olfactory bulbs and epithelium were harvested from an independent cohort of female hamsters at 24 dpi and assessed for the presence of **(F)** *Isg15* and **(G)** *Ccl5* transcripts via qRT-PCR.

**(H-J)** The buried food-finding test was performed on hamsters at (**H)** 3, **(I)** 15, and **(J)** 28dpi. Following testing with time measured to discovery of buried food, the test was also repeated with food that was visible rather than buried.

## METHODS

### Data visualization

All non-RNA-Seq statistical analyses, box and bar graphs, and Kaplan-Meyer plots were prepared using prism 9 as described in figure legends (GraphPad Software, San Diego, California USA; https://www.graphpad.com/).

### Virus and cells

SARS-CoV-2 isolate USA-WA1/2020 was propagated in Vero-E6 cells in DMEM supplemented with 2% FBS, 1mM HEPES and 1% penicillin/streptomycin. Virus stocks were filtered via centrifugation with Amicon Ultra-15 Centrifugal filter unit (Sigma). Infectious viral titers were quantified by plaque assay in Vero-E6 cells in MEM supplemented with 2% FBS, 1mM HEPES and 0.7% OXOID agar (Thermo Fisher). Assays were fixed in 5% paraformaldehyde and stained with crystal violet. All infections were performed with either passage 3 or 4 SARS-CoV-2. Influenza A virus H1N1 isolate A/California/04/2009 was propagated in MDCK cells in DMEM supplemented with 0.35% BSA. All cells were tested for the presence of mycoplasma using MycoAlert Mycoplasma Detection Kit (Lonza).

### Hamster experiments

6–7 week-old male Golden Syrian hamsters (*Mesocricetus auratus*) were obtained from Charles River Laboratories. Hamsters were acclimated to the CDC/USDA-approved BSL-3 facility at the Icahn School of Medicine at Mount Sinai or NYUL for at least seven days. Hamsters were intranasally infected with PBS, 1000 PFU (100uL) of SARS-CoV-2, or 100,000 PFU (100 uL) of H1N1 influenza A virus under ketamine/xylazine anesthesia. Hamsters were euthanized via intraperitoneal injection of pentobarbital and cardiac perfusion with 60 mL PBS. Each tissue was harvested at day 1-, 3-, 4-, 7-, 14-, or 31-days post-infection. Collected tissues were homogenized with PBS or Trizol (Thermo Fisher) in Lysing Matrix A homogenization tubes (MP Biomedicals) for 40 seconds at 6 m/s for 2 cycles in a FastPrep 24 5G bead grinder and lysis system (MP Biomedicals) for plaque assay or RNA isolation, respectively. Additional tissues were fixed in 4% paraformaldehyde for >72 hours prior to embedding in paraffin wax blocks for histology. Prior to fixation, lungs were inflated using 1.5 mL of 4% PFA administered via intratracheal catheter. An independent female cohort of 6-7 week-old female Golden Syrian hamsters was also obtained from Charles River Laboratories and treated in an analogous manner. These hamsters were sacrificed at 24-days-post-infection, and collected tissues were processed in an identical manner. All animal experiments were performed according to protocols approved by the Institutional Animal Care and Use Committee (IACUC) and Institutional Biosafety Committee at ISMMS and NYUL. Hamsters were randomly assigned to the different treatment groups and all IAV and SARS-CoV-2 infections were performed in the BSL-3 facility.

### qRT-PCR

RNA was isolated from homogenized samples by TRIzol/phenol-chloroform extraction. 1ug of total RNA from each tissue was reverse-transcribed into cDNA with oligo dT primers using SuperScript II Reverse transcriptase (Thermo Fisher). Quantitative RT-PCR was performed using primers described in table S2 and KAPA SYBR Fast qPCR Master Mix (KAPA Biosystems) on a LightCycler 480 Instrument II (Roche). Delta-delta-cycle threshold (DDCt) was determined relative to mock-infected control unless otherwise stated (Hoagland et al., 2021).

### Hamster RNA sequencing

RNA was isolated from homogenized samples by TRIzol/phenol-chloroform extraction. 1 ug of total RNA from each tissue was enriched for polyadenylated RNA and prepared for next-generation sequencing using the TruSeq Stranded mRNA Library Prep Kit (Illumina) according to manufacturer’s instructions. Prepared libraries were sequenced on an Illumina NextSeq 500 platform. Fastq files were generated with bcl2fastq (Illumina) and aligned to the Syrian golden hamster genome (MesAur 1.0, ensembl) using the RNA-Seq Alignment application (Basespace, Illumina). Salmon files were analyzed using the DESeq2 analysis pipeline (Love et al., 2014). All genes with an adjusted p value (p-adj) of <0.1 were considered Differentially Expressed Genes (DEGs). Gene Set Enrichment Analysis (GSEA) studies were performed using the GSEA_4.1.0 Mac App as made available by the Broad Institute and UC San Diego (Mootha et al., 2003; Subramanian et al., 2005). Analyses were conducted on a pre-ranked gene list with ranking statistic calculated from DESeq2 results output as follows: -log10(p-value)*sign(log2FoldChange) (Reimand et al., 2019). Unbiased GSEA analyses were conducted against the Hallmark Gene Sets (v7.4), the curated C5 gene ontology and human phenotype ontology gene set (v7.4), and the curated C8 cell type signature gene sets (v7.4) made available by the Molecular Signatures Database (MSigDB). Additional GSEA analyses were conducted on gene sets manually curated from prior publications as described in text. All visualizations of RNA-sequencing differential expression data were created in R using ggplot2, pheatmap, ComplexHeatmap, and gplots packages. Gene set enrichment plots were adapted from VisualizeRNAseq (https://github.com/GryderArt/VisualizeRNAseq). Radar plots were created using the ggradar2 package (https://github.com/xl0418/ggradar2). Assessment of read coverage of viral genome was conducted using Bowtie2 and IGV_2.8.13 and visualized using ggplot2. Rank-rank scatter plots were created using the RRHO package using the same ranking statistic as was used in GSEA analyses (Plaisier et al., 2010).

### H&E, Verhoeff Van Gieson, TUNEL Staining and Quantification

Paraffin-embedded tissue blocks were cut into 5-micron sections and mounted on charged glass slides. Sections were deparaffinized by immersion in xylene and rehydrated in decreasing ethanol dilutions. Slides were then stained with hematoxylin (Gill’s formula, Vector Laboratories, Cat #H3401) and eosin Y (Sigma Aldrich, Cat #E4009) according to manufacturer’s instructions. Slides were dehydrated via immersion in increasing concentrations of ethanol, cleared with xylene, and coverslipped (Hoagland et al., 2021). Sections were assessed for clinical features by a board-certified pathologist. Images were morphometrically analyzed using QuPath (Bankhead et al., 2017) and ImageJ (Schneider et al., 2012). Randomly sampled tissue regions were generated from digitized lung and kidney histological images. In kidneys, these regions were assessed for average cellular size across each area. In lung sampled areas, lambertosis coverage and airway sizes were manually quantified by treatment-blinded team members. Verhoeff Van Gieson staining was performed on 5-micron sections that were cut from paraffin-embedded tissue blocks and embedded on charged glass slides. Slides were stained using ‘Elasic Stain Kit (Verhoeff Van Gieson/EVG Stain)’ kit (Abcam, ab150667) according to manufacturer instructions. Slides were dehydrated via immersion in increasing concentrations of ethanol, cleared with xylene, and coverslipped (Hoagland et al., 2021). Slides were digitized using Hamamatsu S210 digital slide scanner. All images of slides were captured using NDP.view.2 software (Hamamatsu).

TUNEL staining was performed on 5-micron sections that were cut from paraffin-embedded tissue blocks and embedded on charged glass slides. Slides were deparaffinized and processed using the ‘TUNEL Assay Kit –BrdU-Red’ kit (Abcam, ab666110) according to manufacturer instructions. Nuclei were additionally stained with DAPI. Slides were coverslipped and assessed via confocal microscopy. Total number of TUNEL positive nuclei per cross-section of tissue were manually tabulated.

### Immunohistochemistry

Paraffin-embedded tissue blocks were cut into 5-micron sections and mounted on charged glass slides. Sections were deparaffinized by immersion in xylene and rehydrated in decreasing ethanol dilutions. Antigen retrieval was performed for 45 minutes in IHC-Tek Epitope Retrieval Steamer (Cat #IW-1102) with slides immersed in IHC-Tek Epitope Retrieval Solution (Cat #IW-1100). Tissue was blocked in TBS with 10% goat serum and 1% bovine serum albumin. Primary antibody (MX-A: Millipore Sigma, MABF938; IBA-1: Wako, 019-19741; CD3: Dako, A0452; MPO: Dako, A0398) was added to slides at a dilution (MX-A, 1:100; IBA-1, 1:1000; CD3, 1:1000; MPO: 1:5000), and sections were incubated overnight at 4 degrees Celsius. Slides were washed in TBS with 0.025% Triton X-100 prior to immersion in 0.3% hydrogen peroxide in TBS for 15 minutes. Slides were washed once again and HRP-conjugated secondary antibody was added at a 1:5000 concentration (Goat anti-mouse: ThermoFisher, Cat #A21426; Goat anti-rabbit: Abcam, Ab6721). Slides were washed twice prior to application of DAB developing reagent (Vector Laboratories, Cat #SK-4105). Slides were dehydrated via immersion in increasing concentrations of ethanol, cleared with xylene, and coverslipped. Slides were digitized using Hamamatsu S210 digital slide scanner. All images of slides were captured using NDP.view.2 software (Hamamatsu).

### Olfaction Assessment

Olfaction was assessed via the buried food finding test as previously described (de Melo *et al*., 2021; Lazarini et al., 2018). Hamsters were presented with cereal (Coco Krispies, Kellogs) five days prior to test; all were consumed within 1 hour. 20 hours prior to testing, hamsters were food restricted. On the day of testing, hamsters were placed into clean cages with standard bedding and allowed to acclimate for 20 minutes. After 20 minutes, hamsters were moved to a holding cage for two minutes while chocolate cereal was buried underneath the bedding in a corner of the testing cage. Hamsters were then moved back to the cage with cereal in it and placed in the opposite corner of the cage as the buried cereal. Hamsters were timed from placement in cage to the time of detection of food (digging in the area of the buried cereal). Hamsters were limited to a 15-minute maximum period to find cereal. Once food was found, hamsters were moved back to holding cage for one minute, and food was placed on top of bedding (visible) in a corner of the test cage during this time. The hamster was then once again placed in the opposite corner of the test cage from the cereal, and time was recorded from placement of hamster in cage to detection of food. All behavioral studies were in compliance with institutional IACUC protocols and took place inside of a biosafety cabinet according to BSL-3 protocols.

### Marble Burying Assay

The marble burying assay was adapted from previously described protocols (Yanai and Endo, 2021). Hamsters were placed into a corner of a cage with clean bedding that had 20 equally-spaced glass marbles placed inside of it. Hamsters were allowed to move freely about the cage for 15 minutes, at which time they were moved back to their original cage. The number of buried and unburied marbles per cage were tallied by two independent observers and averaged. Partially buried marbles were counted as buried if greater than 60% of the marble was covered with bedding material. All group were assessed for outliers which were corrected for using Iterative Grubb’s method. All behavioral studies were in compliance with institutional IACUC protocols and took place inside of a biosafety cabinet according to BSL-3 protocols.

### RNA fluorescent in-situ hybridization (RNAscope)

The Fluorescent Multiplex V2 kit (Advanced Cell Diagnostics, CA) was used for RNAscope FISH. Specifically, we used the FFPE protocol as detailed in the RNAscope Multiplex Fluorescent Reagent Kit v2 Assay User Manual. RNAscope probes were as follows: *Rbfox3* (NeuN) for pan-neuronal labeling (Mau-Rbfox3-C1) and the Spike gene (*S*) for SARS-CoV-2 labeling (V-nCoV2019-S-C3). Opal dyes (Akoya Biosciences, MA) were used for secondary staining as follows: Opal 690 for C1 and Opal 570 for C3. DAPI was used for nuclear staining. Images were taken on an LSM880 confocal microscope (Zeiss, GER) with identical parameters between mock- and SARS-CoV-2-infected samples.

### IRB Statement

Tissue human samples were provided by the Weill Cornell Medicine Department of Pathology. The Tissue Procurement Facility operates under Institutional Review Board (IRB) approved protocol and follows guidelines set by Health Insurance Portability and Accountability Act (HIPAA). Experiments using samples from human subjects were conducted in accordance with local regulations and with the approval of the IRB at the Weill Cornell Medicine. The autopsy samples are considered human tissue research and were collected under IRB protocols 20-04021814 and 19-11021069. All autopsies have consent for research use from next of kin, and these studies were determined as exempt by IRB at Weill Cornell Medicine under those protocol numbers.

### Heart, Lung, Kidney Patient sample collection

All autopsies are performed with consent of next of kin and permission for retention and research use of tissue. Autopsies were performed in a negative pressure room with protective equipment including N-95 masks; brain and bone were not obtained for safety reasons. All fresh tissues were procured prior to fixation and directly into Trizol for downstream RNA extraction. Tissues were collected from lung, kidney, and the heart as consent permitted. Post-mortem intervals ranged from less than 24 hours to 72 hours (with 2 exceptions - one at 4 and one at 7 days - but passing RNA quality metrics) with an average of 2.5 days. All deceased patient remains were refrigerated at 4°C prior to autopsy performance.

### Human Heart, Lung, Kidney RNA-sequencing

For RNA library preparation, all samples’ RNA was treated with DNAse 1 (Zymo Research, Catalog # E1010). Post-DNAse digested samples were then put into the NEBNext rRNA depletion v2 (Human/Mouse/Rat), Ultra II Directional RNA (10ng), and Unique Dual Index Primer Pairs were used following the vendor protocols from New England Biolabs. Completed libraries were quantified by Qubit and run on a Bioanalyzer for size determination. Libraries were pooled and sent to the WCM Genomics Core or HudsonAlpha for final quantification by Qubit fluorometer (ThermoFisher Scientific), TapeStation 2200 (Agilent), and qRT-PCR using the Kapa Biosystems Illumina library quantification kit.

NYGC RNA sequencing libraries were prepared using the KAPA Hyper Library Preparation Kit + RiboErase, HMR (Roche) in accordance with manufacturer’s recommendations. Briefly, 50-200ng of Total RNA were used for ribosomal depletion and fragmentation. Depleted RNA underwent first and second strand cDNA synthesis followed by adenylation, and ligation of unique dual indexed adapters. Libraries were amplified using 12 cycles of PCR and cleaned-up by magnetic bead purification. Final libraries were quantified using fluorescent-based assays including PicoGreen (Life Technologies) or Qubit Fluorometer (Invitrogen) and Fragment Analyzer (Advanced Analytics) and sequenced on a NovaSeq 6000 sequencer (v1 chemistry) with 2×150bp targeting 60M reads per sample.

RNAseq data was processed through the nf-core/rnaseq pipeline (Ewels et al., 2020). This workflow involved quality control of the reads with FastQC (Andrews), adapter trimming using Trim Galore! (https://github.com/FelixKrueger/TrimGalore), read alignment with STAR (Dobin et al., 2013), gene quantification with Salmon (Patro et al., 2017), duplicate read marking with Picard MarkDuplicates (https://github.com/broadinstitute/picard), and transcript quantification with StringTie (Kovaka et al., 2019). Other quality control measures included RSeQC, Qualimap, and dupRadar. Alignment was performed using the GRCh38 build native to nf-core and annotation was performed using Gencode Human Release 33 (GRCH38.p13). Differential expression of genes was calculated by DESeq2 using FeatureCounts reads. Differential expression comparisons were done as either COVID high cases versus COVID-controls or COVID low cases versus COVID-controls for each tissue specifically. COVID viral load designations were assigned after quantification of normalized reads mapping to the SARS-CoV-2 genome for each donor. Genes were ranked by the following statistic: log10(p-value)*sign(log2FoldChange). Ranked genes were used as input for gene set enrichment analysis (GSEA) on the molecular signatures database (MSigDB).

### Human Olfactory Bulb and Olfactory Epithelium Sequencing

Brain tissue and nasal epithelium, including the olfactory epithelium and bulb were retrieved under a collaborative effort by the Department of Neuropathology and the Department of Otolaryngology at Columbia University Irving Medical Center (New York, NY, USA). The study was approved by the ethics and Institutional Review Board of Columbia University Medical Center (IRB AAAT0689, AAAS7370). Nasal tissues, including olfactory and respiratory epithelium were harvested from the skull base using an en-bloc resection of the anterior skull base including the cribriform plate. Olfactory epithelium was isolated from the olfactory cleft, spanning turbinate and adjacent septal mucosa prior to being preserved in Trizol reagent.

For human OE and OB RNA was extracted from 10mg of tissue per sample using Direct-zol RNA kit from Zymo Research (Catalog #R2052). After DNAse treatment 50ng-1ug of total RNA was used to prepare DNA libraries with Truseq RNA Library Prep Kit v2 (Illumina) following manufacture’s instruction. Libraries were amplified using 14 PCR cycles followed by AMPure XP beads purification. Next, libraries were quantified with Bioanalyser (Agilent Technologies) and Qubit (Invitrogen) and sequenced on NovaSeq 6000 sequencer (Illumina) at Columbia Genome Center.

All resulting fastq files were aligned to the Homo Sapiens genome (GRCh38, RefSeq) using the RNA-Seq Alignment application (Basespace, Illumina). Salmon files were analyzed using the DESeq2 analysis pipeline (Love et al., 2014). All genes with an adjusted p value (p-adj) of <0.1 were considered Differentially Expressed Genes (DEGs). Gene Set Enrichment Analysis (GSEA) studies were performed as described above in ‘Hamster RNA Sequencing’.

### Graphic Creation

All graphics were created using BioRender and Microsoft Powerpoint.

