## Supplementary figures and images for "SARS-CoV-2 infection results in lasting and systemic perturbations post recovery"

### Supplemental Figures

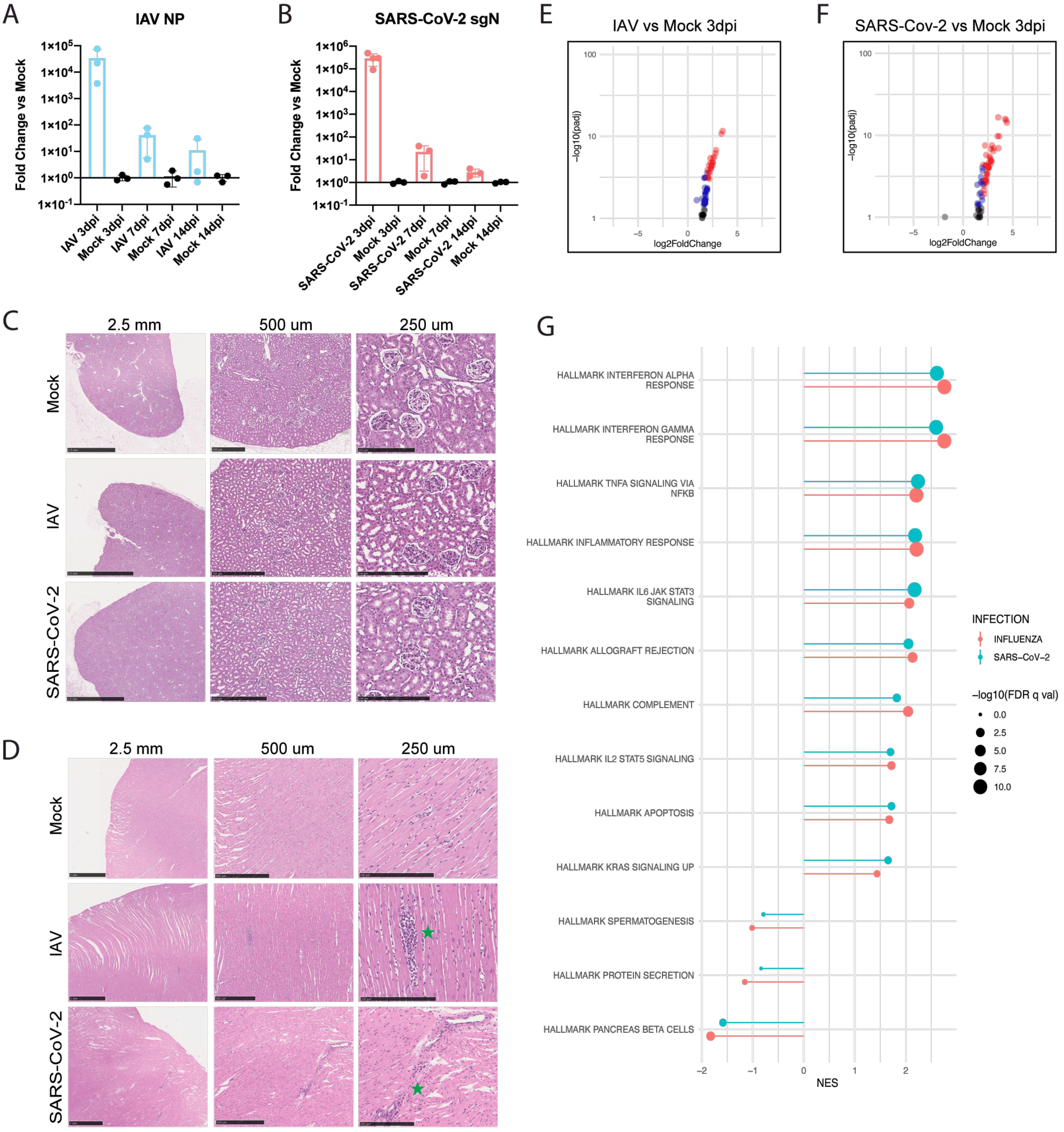

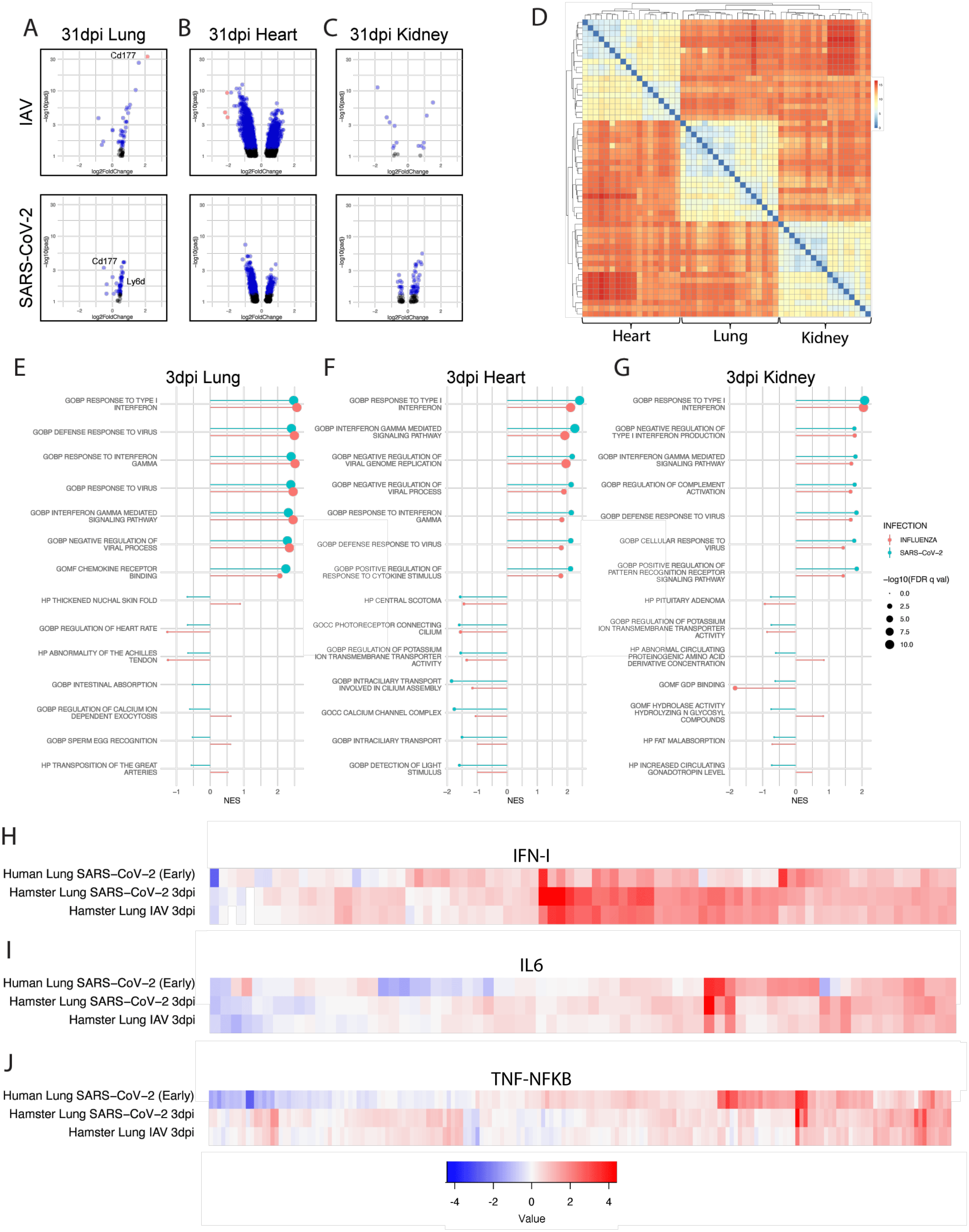

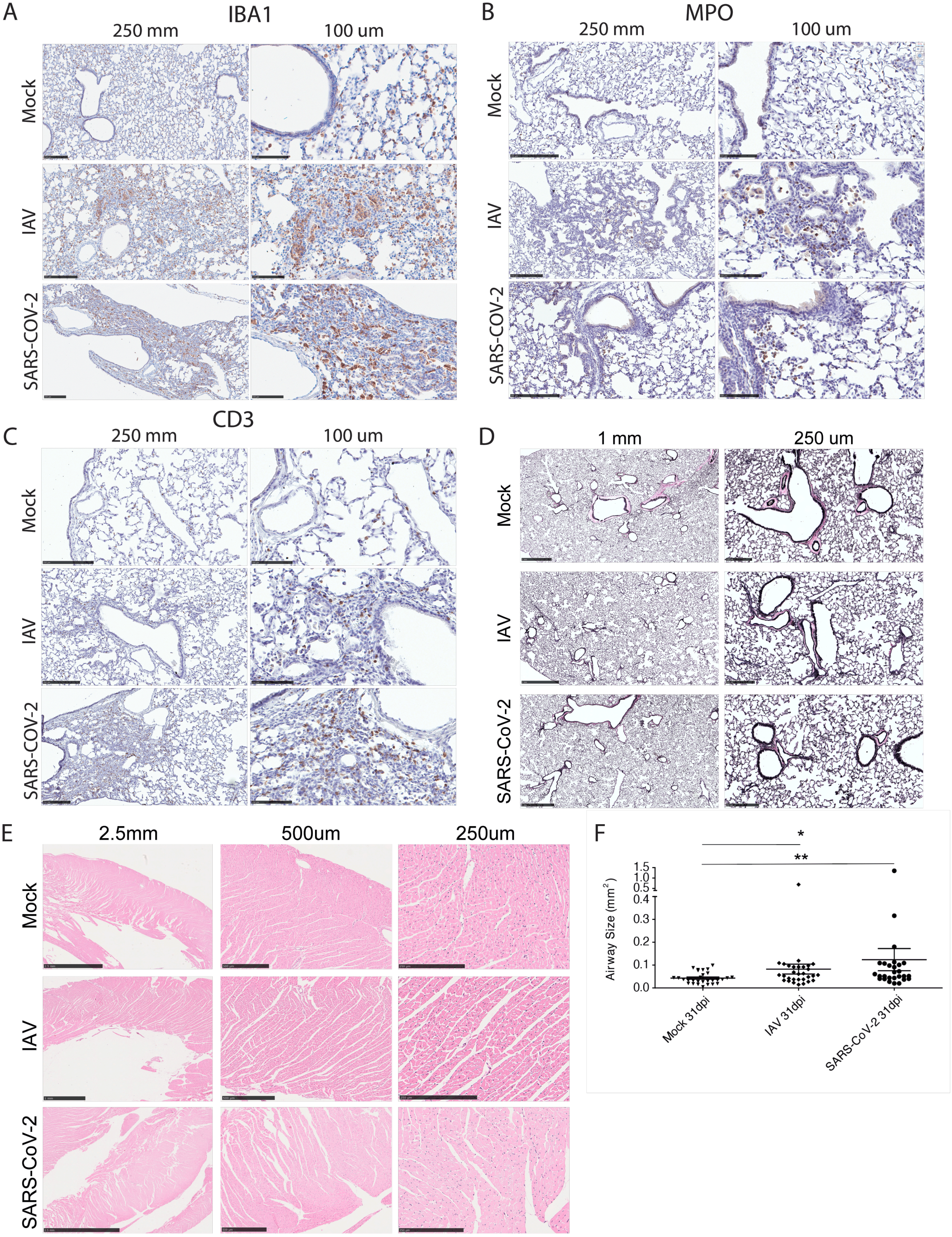

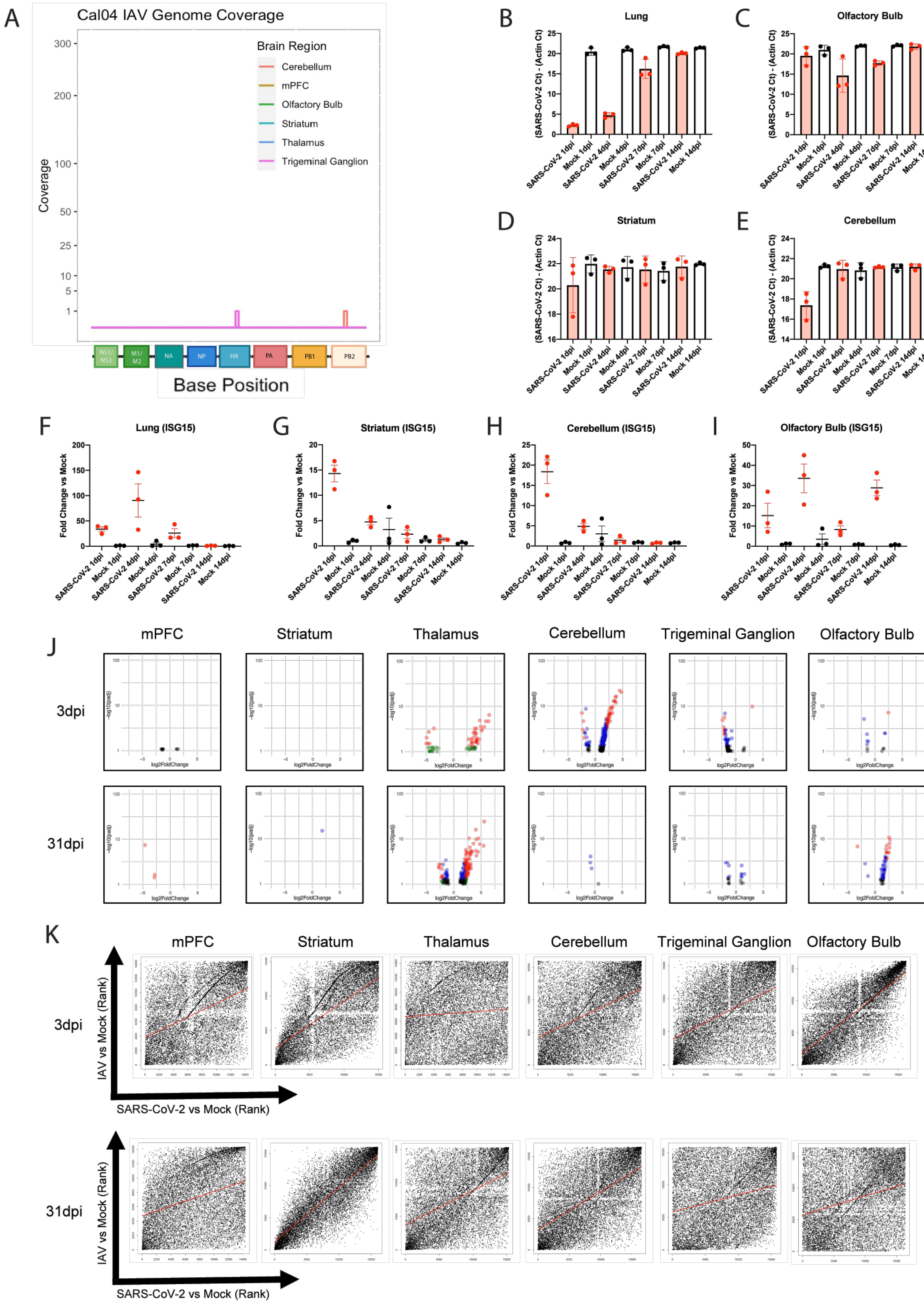

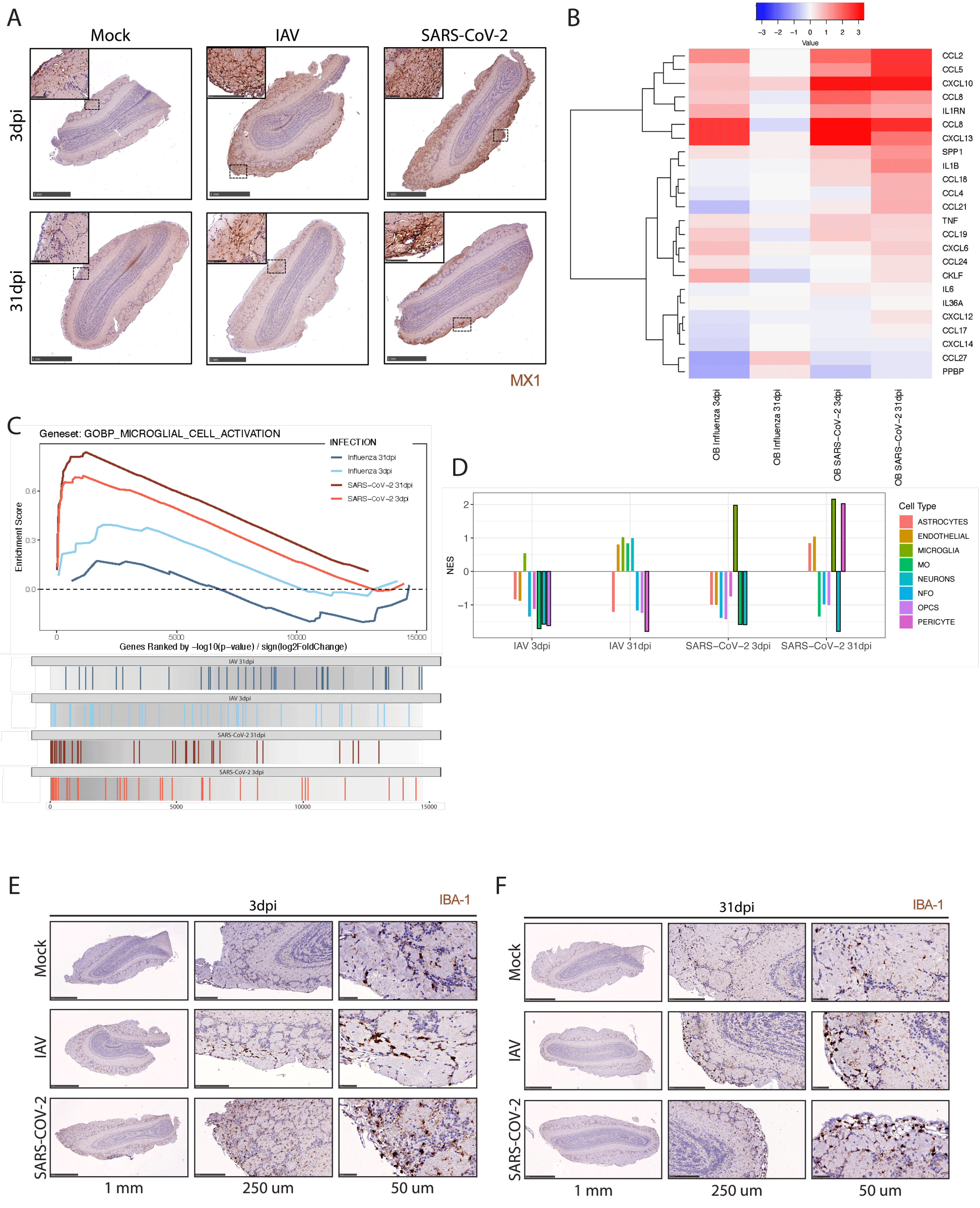

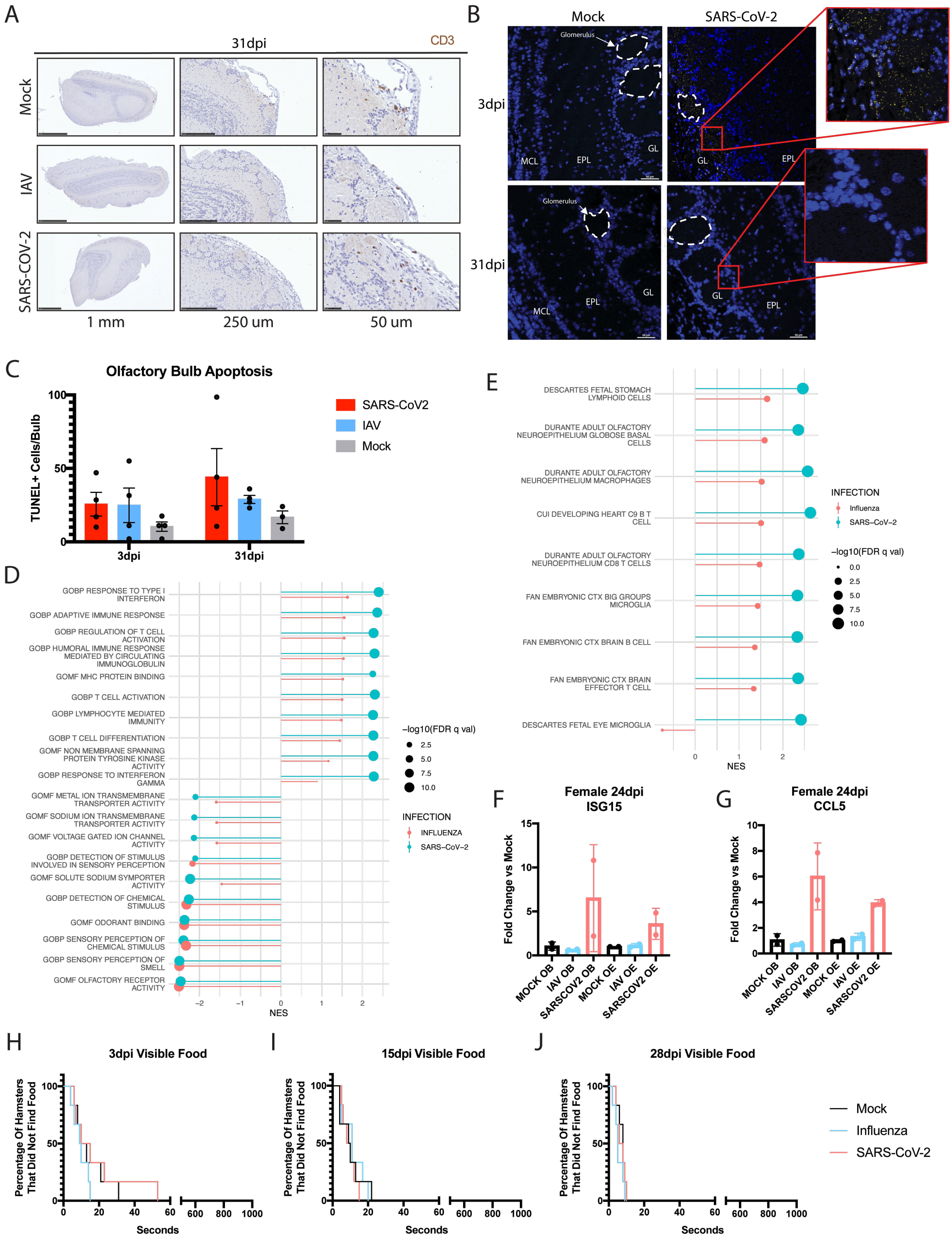
