## Supplemental Tables for "SARS-CoV-2 infection results in lasting and systemic perturbations post recovery"

### Table S1: Differentially Expressed Genes (p-adj < 0.1) for 3dpi Lung Infection vs. Mock

| Comparison | Ensembl Gene ID | External Gene Name | Base Mean | log2 Fold Change | lfcSE | stat | pvalue | padj |
|---|---|---|---|---|---|---|---|---|
| **SARS-CoV-2 vs. Mock (3dpi lung)** | ENSMAUG00000000088 | Mx1 | 267.422303 | 2.80263335 | 0.44785895 | 6.32640816 | 2.51E-10 | 5.52E-07 |
| **SARS-CoV-2 vs. Mock (3dpi lung)** | ENSMAUG00000000309 | Eif2ak2 | 211.415543 | 1.6729575 | 0.44332446 | 3.77891328 | 0.00015751 | 0.02950544 |
| **SARS-CoV-2 vs. Mock (3dpi lung)** | ENSMAUG00000000781 | Ccl5 | 33.753774 | 1.76244833 | 0.47477966 | 3.8538927 | 0.00011625 | 0.02436917 |
| **SARS-CoV-2 vs. Mock (3dpi lung)** | ENSMAUG00000000842 | Ccr5 | 75.49706 | 1.74716226 | 0.46742217 | 3.80652415 | 0.00014093 | 0.0288192 |
| **SARS-CoV-2 vs. Mock (3dpi lung)** | ENSMAUG00000000857 | Helz2 | 216.228036 | 1.65836771 | 0.35031758 | 4.73574697 | 2.18E-06 | 0.00076859 |
| **SARS-CoV-2 vs. Mock (3dpi lung)** | ENSMAUG00000000936 | | 91.2628387 | 2.06839873 | 0.43562883 | 4.75463333 | 1.99E-06 | 0.00072929 |
| **SARS-CoV-2 vs. Mock (3dpi lung)** | ENSMAUG00000002308 | Calhm6 | 103.317359 | 1.76981781 | 0.47316023 | 3.9137973 | 9.09E-05 | 0.02051014 |
| **SARS-CoV-2 vs. Mock (3dpi lung)** | ENSMAUG00000002531 | Slamf9 | 63.6555647 | 0.91283403 | 0.40903152 | 3.89527341 | 9.81E-05 | 0.02158917 |
| **SARS-CoV-2 vs. Mock (3dpi lung)** | ENSMAUG00000002664 | Parp9 | 97.7211953 | 1.79848173 | 0.42751899 | 4.18616806 | 2.84E-05 | 0.00780539 |
| **SARS-CoV-2 vs. Mock (3dpi lung)** | ENSMAUG00000003013 | Slfn1 | 14.4635636 | 1.54660529 | 0.47294874 | 3.46825674 | 0.00052385 | 0.07911325 |
| **SARS-CoV-2 vs. Mock (3dpi lung)** | ENSMAUG00000003378 | Hk1 | 115.744923 | 2.3045124 | 0.38967421 | 5.93371352 | 2.96E-09 | 3.72E-06 |
| **SARS-CoV-2 vs. Mock (3dpi lung)** | ENSMAUG00000003396 | Psmb8 | 37.1017713 | 1.54793288 | 0.45535336 | 3.48315946 | 0.00049553 | 0.07879778 |
| **SARS-CoV-2 vs. Mock (3dpi lung)** | ENSMAUG00000004249 | | 96.3064014 | 1.73124641 | 0.46464803 | 3.76122564 | 0.00016908 | 0.0309167 |
| **SARS-CoV-2 vs. Mock (3dpi lung)** | ENSMAUG00000004294 | | 788.93254 | 3.51687274 | 0.45843046 | 8.1854256 | 2.71E-16 | 2.39E-12 |
| **SARS-CoV-2 vs. Mock (3dpi lung)** | ENSMAUG00000004507 | Ifit2 | 71.7251356 | 2.34010598 | 0.46218509 | 5.25814475 | 1.46E-07 | 8.54E-05 |
| **SARS-CoV-2 vs. Mock (3dpi lung)** | ENSMAUG00000005083 | | 227.348063 | 1.62101879 | 0.4397455 | 3.72286464 | 0.00019698 | 0.03468338 |
| **SARS-CoV-2 vs. Mock (3dpi lung)** | ENSMAUG00000005220 | Ifi35 | 81.5714862 | 1.70549855 | 0.45409843 | 3.79275824 | 0.00014898 | 0.0288192 |
| **SARS-CoV-2 vs. Mock (3dpi lung)** | ENSMAUG00000006907 | | 32.9796824 | 1.38685295 | 0.46623236 | 3.48076478 | 0.00049998 | 0.07879778 |
| **SARS-CoV-2 vs. Mock (3dpi lung)** | ENSMAUG00000007175 | | 269.999158 | 1.76502907 | 0.37013234 | 4.76044849 | 1.93E-06 | 0.00072929 |
| **SARS-CoV-2 vs. Mock (3dpi lung)** | ENSMAUG00000007782 | | 224.515717 | 2.37905461 | 0.43423527 | 5.45439728 | 4.91E-08 | 3.61E-05 |
| **SARS-CoV-2 vs. Mock (3dpi lung)** | ENSMAUG00000007959 | H2-T24 | 23.979144 | 1.70945855 | 0.44964956 | 3.79477767 | 0.00014778 | 0.0288192 |
| **SARS-CoV-2 vs. Mock (3dpi lung)** | ENSMAUG00000008273 | AA467197 | 93.7183089 | 1.96472123 | 0.47316554 | 5.00465151 | 5.60E-07 | 0.00024635 |
| **SARS-CoV-2 vs. Mock (3dpi lung)** | ENSMAUG00000008425 | | 34.9942246 | 2.17707594 | 0.45973783 | 4.69793885 | 2.63E-06 | 0.00088988 |
| **SARS-CoV-2 vs. Mock (3dpi lung)** | ENSMAUG00000008669 | Rsad2 | 51.5212698 | 2.31359305 | 0.47388737 | 5.63538209 | 1.75E-08 | 1.54E-05 |
| **SARS-CoV-2 vs. Mock (3dpi lung)** | ENSMAUG00000009426 | Isg15 | 93.8637336 | 2.84433679 | 0.47222855 | 6.55641194 | 5.51E-11 | 1.62E-07 |
| **SARS-CoV-2 vs. Mock (3dpi lung)** | ENSMAUG00000009739 | | 75.5775456 | 2.37514769 | 0.46533304 | 5.3671338 | 8.00E-08 | 5.03E-05 |
| **SARS-CoV-2 vs. Mock (3dpi lung)** | ENSMAUG00000009900 | | 41.0934167 | 1.78835736 | 0.4681336 | 4.08365987 | 4.43E-05 | 0.01115137 |
| **SARS-CoV-2 vs. Mock (3dpi lung)** | ENSMAUG00000010500 | | 107.88939 | 1.93318592 | 0.45533384 | 4.26594576 | 1.99E-05 | 0.00584166 |
| **SARS-CoV-2 vs. Mock (3dpi lung)** | ENSMAUG00000010928 | Gbp5 | 175.721725 | 1.73215877 | 0.39526143 | 4.35793535 | 1.31E-05 | 0.00428119 |
| **SARS-CoV-2 vs. Mock (3dpi lung)** | ENSMAUG00000010948 | | 62.5091322 | 2.56985422 | 0.469514 | 5.95139105 | 2.66E-09 | 3.72E-06 |
| **SARS-CoV-2 vs. Mock (3dpi lung)** | ENSMAUG00000011314 | Ccl7 | 44.1220415 | 2.01658273 | 0.47266129 | 5.15802685 | 2.50E-07 | 0.00012925 |
| **SARS-CoV-2 vs. Mock (3dpi lung)** | ENSMAUG00000012413 | Ifi44 | 228.344592 | 2.11702848 | 0.4219789 | 5.00791474 | 5.50E-07 | 0.00024635 |
| **SARS-CoV-2 vs. Mock (3dpi lung)** | ENSMAUG00000013430 | | 39.433565 | 1.8179204 | 0.4671428 | 3.97116108 | 7.15E-05 | 0.0174914 |
| **SARS-CoV-2 vs. Mock (3dpi lung)** | ENSMAUG00000013633 | Plaat3 | 83.0754512 | 1.44661853 | 0.3650809 | 3.95047642 | 7.80E-05 | 0.01855878 |
| **SARS-CoV-2 vs. Mock (3dpi lung)** | ENSMAUG00000014213 | Cxcl10 | 76.2127146 | 2.15921182 | 0.47384771 | 5.43015409 | 5.63E-08 | 3.81E-05 |
| **SARS-CoV-2 vs. Mock (3dpi lung)** | ENSMAUG00000014262 | Lgals3bp | 90.5492576 | 1.53179085 | 0.46866149 | 3.45337164 | 0.00055363 | 0.08123535 |
| **SARS-CoV-2 vs. Mock (3dpi lung)** | ENSMAUG00000014297 | Arg1 | 153.941858 | 1.95494348 | 0.47220346 | 4.3022478 | 1.69E-05 | 0.00513286 |
| **SARS-CoV-2 vs. Mock (3dpi lung)** | ENSMAUG00000014385 | | 519.20078 | 2.5211303 | 0.450992 | 5.66836158 | 1.44E-08 | 1.54E-05 |
| **SARS-CoV-2 vs. Mock (3dpi lung)** | ENSMAUG00000014482 | Gbp2 | 516.348994 | 2.22020901 | 0.44501028 | 5.02848215 | 4.94E-07 | 0.00024181 |
| **SARS-CoV-2 vs. Mock (3dpi lung)** | ENSMAUG00000014847 | Herc6 | 116.848008 | 1.97230496 | 0.45998722 | 4.33509446 | 1.46E-05 | 0.00458115 |
| **SARS-CoV-2 vs. Mock (3dpi lung)** | ENSMAUG00000015073 | | 9.24856206 | 1.46687202 | 0.465342 | 3.50405034 | 0.00045824 | 0.0761195 |
| **SARS-CoV-2 vs. Mock (3dpi lung)** | ENSMAUG00000015796 | Adar | 119.341852 | 1.74603062 | 0.36392156 | 4.7711515 | 1.83E-06 | 0.00072929 |
| **SARS-CoV-2 vs. Mock (3dpi lung)** | ENSMAUG00000016555 | Xdh | 673.123079 | 1.4536058 | 0.42814636 | 3.39863956 | 0.00067722 | 0.09768484 |
| **SARS-CoV-2 vs. Mock (3dpi lung)** | ENSMAUG00000016558 | Ube2l6 | 20.2841472 | 1.6562272 | 0.47425838 | 3.88208358 | 0.00010357 | 0.02223875 |
| **SARS-CoV-2 vs. Mock (3dpi lung)** | ENSMAUG00000016664 | Ddx60 | 103.677813 | 1.77594725 | 0.43444402 | 4.09156205 | 4.28E-05 | 0.01109504 |
| **SARS-CoV-2 vs. Mock (3dpi lung)** | ENSMAUG00000016788 | Gar1 | 53.8052528 | 1.81124174 | 0.47043052 | 3.91953788 | 8.87E-05 | 0.02051014 |
| **SARS-CoV-2 vs. Mock (3dpi lung)** | ENSMAUG00000016905 | | 122.800512 | 2.49289446 | 0.45832613 | 5.46000023 | 4.76E-08 | 3.61E-05 |
| **SARS-CoV-2 vs. Mock (3dpi lung)** | ENSMAUG00000016913 | Mx2 | 1031.78846 | 3.39625503 | 0.45846043 | 7.8502 | 4.15E-15 | 1.83E-11 |
| **SARS-CoV-2 vs. Mock (3dpi lung)** | ENSMAUG00000018005 | | 62.1010941 | 1.46594038 | 0.43516913 | 3.39434831 | 0.00068792 | 0.09768484 |
| **SARS-CoV-2 vs. Mock (3dpi lung)** | ENSMAUG00000018176 | Ccl8 | 148.658814 | 1.66805262 | 0.46913296 | 3.61859539 | 0.00029621 | 0.05113334 |
| **SARS-CoV-2 vs. Mock (3dpi lung)** | ENSMAUG00000018346 | Gbp2b | 46.6527113 | 1.96182149 | 0.47244659 | 4.23477824 | 2.29E-05 | 0.00649728 |
| **SARS-CoV-2 vs. Mock (3dpi lung)** | ENSMAUG00000018871 | Fcgr4 | 33.3494471 | 1.68081568 | 0.47455148 | 3.75684119 | 0.00017207 | 0.0309167 |
| **SARS-CoV-2 vs. Mock (3dpi lung)** | ENSMAUG00000018991 | | 51.523171 | 1.61345546 | 0.46448612 | 3.48010751 | 0.00050121 | 0.07879778 |
| **SARS-CoV-2 vs. Mock (3dpi lung)** | ENSMAUG00000019294 | Hp | 54.3518025 | 1.59254979 | 0.47044832 | 3.46502743 | 0.00053018 | 0.07911325 |
| **SARS-CoV-2 vs. Mock (3dpi lung)** | ENSMAUG00000019385 | Rnf114 | 158.113477 | 1.44786746 | 0.41560323 | 3.465525 | 0.0005292 | 0.07911325 |
| **SARS-CoV-2 vs. Mock (3dpi lung)** | ENSMAUG00000019671 | Gbp7 | 40.0982502 | 1.77586149 | 0.46706397 | 3.79011526 | 0.00015058 | 0.0288192 |
| **SARS-CoV-2 vs. Mock (3dpi lung)** | ENSMAUG00000019884 | Irf7 | 51.8339066 | 2.51972448 | 0.47350521 | 5.6467489 | 1.64E-08 | 1.54E-05 |
| **SARS-CoV-2 vs. Mock (3dpi lung)** | ENSMAUG00000019934 | Dhx58 | 28.09127 | 1.69010452 | 0.47437136 | 3.58628469 | 0.00033542 | 0.05678966 |
| **SARS-CoV-2 vs. Mock (3dpi lung)** | ENSMAUG00000020370 | Irf9 | 68.1029628 | 1.75783614 | 0.42269541 | 4.15389544 | 3.27E-05 | 0.0087203 |
| **SARS-CoV-2 vs. Mock (3dpi lung)** | ENSMAUG00000021773 | Uba7 | 36.0443047 | 2.32691239 | 0.44735307 | 5.2339316 | 1.66E-07 | 9.13E-05 |
| **SARS-CoV-2 vs. Mock (3dpi lung)** | ENSMAUG00000021814 | Ddx58 | 160.376496 | 2.02712186 | 0.42366247 | 4.77051739 | 1.84E-06 | 0.00072929 |
| **SARS-CoV-2 vs. Mock (3dpi lung)** | ENSMAUG00000022167 | Apobec1 | 120.186745 | 2.68053078 | 0.46694968 | 6.08059071 | 1.20E-09 | 2.11E-06 |
| **IAV vs. Mock** | ENSMAUG00000000088 | Mx1 | 267.422303 | 2.80263335 | 0.44785895 | 6.32640816 | 2.51E-10 | 5.52E-07 |
| **IAV vs. Mock** | ENSMAUG00000000309 | Eif2ak2 | 211.415543 | 1.6729575 | 0.44332446 | 3.77891328 | 0.00015751 | 0.02950544 |
| **IAV vs. Mock** | ENSMAUG00000000781 | Ccl5 | 33.753774 | 1.76244833 | 0.47477966 | 3.8538927 | 0.00011625 | 0.02436917 |
| **IAV vs. Mock** | ENSMAUG00000000842 | Ccr5 | 75.49706 | 1.74716226 | 0.46742217 | 3.80652415 | 0.00014093 | 0.0288192 |
| **IAV vs. Mock** | ENSMAUG00000000857 | Helz2 | 216.228036 | 1.65836771 | 0.35031758 | 4.73574697 | 2.18E-06 | 0.00076859 |
| **IAV vs. Mock** | ENSMAUG00000000936 | | 91.2628387 | 2.06839873 | 0.43562883 | 4.75463333 | 1.99E-06 | 0.00072929 |
| **IAV vs. Mock** | ENSMAUG00000002308 | Calhm6 | 103.317359 | 1.76981781 | 0.47316023 | 3.9137973 | 9.09E-05 | 0.02051014 |

| | | | | | | | | |
|---|---|---|---|---|---|---|---|---|
| **IAV vs. Mock** | ENSMAUG00000002531 | Slamf9 | 63.6555647 | 0.91283403 | 0.40903152 | 3.89527341 | 9.81E-05 | 0.02158917 |
| **IAV vs. Mock** | ENSMAUG00000002664 | Parp9 | 97.7211953 | 1.79848173 | 0.42751899 | 4.18616806 | 2.84E-05 | 0.00780539 |
| **IAV vs. Mock** | ENSMAUG00000003013 | Slfn1 | 14.4635636 | 1.54660529 | 0.47294874 | 3.46825674 | 0.00052385 | 0.07911325 |
| **IAV vs. Mock** | ENSMAUG00000003378 | Hk1 | 115.744923 | 2.3045124 | 0.38967421 | 5.93371352 | 2.96E-09 | 3.72E-06 |
| **IAV vs. Mock** | ENSMAUG00000003396 | Psmb8 | 37.1017713 | 1.54793288 | 0.45535336 | 3.48315946 | 0.00049553 | 0.07879778 |
| **IAV vs. Mock** | ENSMAUG00000004249 | | 96.3064014 | 1.73124641 | 0.46464803 | 3.76122564 | 0.00016908 | 0.0309167 |
| **IAV vs. Mock** | ENSMAUG00000004294 | | 788.93254 | 3.51687274 | 0.45843046 | 8.1854256 | 2.71E-16 | 2.39E-12 |
| **IAV vs. Mock** | ENSMAUG00000004507 | Ifit2 | 71.7251356 | 2.34010598 | 0.46218509 | 5.25814475 | 1.46E-07 | 8.54E-05 |
| **IAV vs. Mock** | ENSMAUG00000005083 | | 227.348063 | 1.62101879 | 0.4397455 | 3.72286464 | 0.00019698 | 0.03468338 |
| **IAV vs. Mock** | ENSMAUG00000005220 | Ifi35 | 81.5714862 | 1.70549855 | 0.45409843 | 3.79275824 | 0.00014898 | 0.0288192 |
| **IAV vs. Mock** | ENSMAUG00000006907 | | 32.9796824 | 1.38685295 | 0.46623236 | 3.48076478 | 0.00049998 | 0.07879778 |
| **IAV vs. Mock** | ENSMAUG00000007175 | | 269.999158 | 1.76502907 | 0.37013234 | 4.76044849 | 1.93E-06 | 0.00072929 |
| **IAV vs. Mock** | ENSMAUG00000007782 | | 224.515717 | 2.37905461 | 0.43423527 | 5.45439728 | 4.91E-08 | 3.61E-05 |
| **IAV vs. Mock** | ENSMAUG00000007959 | H2-T24 | 23.979144 | 1.70945855 | 0.44964956 | 3.79477767 | 0.00014778 | 0.0288192 |
| **IAV vs. Mock** | ENSMAUG00000008273 | AA467197 | 93.7183089 | 1.96472123 | 0.47316554 | 5.00465151 | 5.60E-07 | 0.00024635 |
| **IAV vs. Mock** | ENSMAUG00000008425 | | 34.9942246 | 2.17707594 | 0.45973783 | 4.69793885 | 2.63E-06 | 0.00088988 |
| **IAV vs. Mock** | ENSMAUG00000008669 | Rsad2 | 51.5212698 | 2.31359305 | 0.47388737 | 5.63538209 | 1.75E-08 | 1.54E-05 |
| **IAV vs. Mock** | ENSMAUG00000009426 | Isg15 | 93.8637336 | 2.84433679 | 0.47222855 | 6.55641194 | 5.51E-11 | 1.62E-07 |
| **IAV vs. Mock** | ENSMAUG00000009739 | | 75.5775456 | 2.37514769 | 0.46533304 | 5.3671338 | 8.00E-08 | 5.03E-05 |
| **IAV vs. Mock** | ENSMAUG00000009900 | | 41.0934167 | 1.78835736 | 0.4681336 | 4.08365987 | 4.43E-05 | 0.01115137 |
| **IAV vs. Mock** | ENSMAUG00000010500 | | 107.88939 | 1.93318592 | 0.45533384 | 4.26594576 | 1.99E-05 | 0.00584166 |
| **IAV vs. Mock** | ENSMAUG00000010928 | Gbp5 | 175.721725 | 1.73215877 | 0.39526143 | 4.35793535 | 1.31E-05 | 0.00428119 |
| **IAV vs. Mock** | ENSMAUG00000010948 | | 62.5091322 | 2.56985422 | 0.469514 | 5.95139105 | 2.66E-09 | 3.72E-06 |
| **IAV vs. Mock** | ENSMAUG00000011314 | Ccl7 | 44.1220415 | 2.01658273 | 0.47266129 | 5.15802685 | 2.50E-07 | 0.00012925 |
| **IAV vs. Mock** | ENSMAUG00000012413 | Ifi44 | 228.344592 | 2.11702848 | 0.4219789 | 5.00791474 | 5.50E-07 | 0.00024635 |
| **IAV vs. Mock** | ENSMAUG00000013430 | | 39.433565 | 1.8179204 | 0.4671428 | 3.97116108 | 7.15E-05 | 0.0174914 |
| **IAV vs. Mock** | ENSMAUG00000013633 | Plaat3 | 83.0754512 | 1.44661853 | 0.3650809 | 3.95047642 | 7.80E-05 | 0.01855878 |
| **IAV vs. Mock** | ENSMAUG00000014213 | Cxcl10 | 76.2127146 | 2.15921182 | 0.47384771 | 5.43015409 | 5.63E-08 | 3.81E-05 |
| **IAV vs. Mock** | ENSMAUG00000014262 | Lgals3bp | 90.5492576 | 1.53179085 | 0.46866149 | 3.45337164 | 0.00055363 | 0.08123535 |
| **IAV vs. Mock** | ENSMAUG00000014297 | Arg1 | 153.941858 | 1.95494348 | 0.47220346 | 4.3022478 | 1.69E-05 | 0.00513286 |
| **IAV vs. Mock** | ENSMAUG00000014385 | | 519.20078 | 2.5211303 | 0.450992 | 5.66836158 | 1.44E-08 | 1.54E-05 |
| **IAV vs. Mock** | ENSMAUG00000014482 | Gbp2 | 516.348994 | 2.22020901 | 0.44501028 | 5.02848215 | 4.94E-07 | 0.00024181 |
| **IAV vs. Mock** | ENSMAUG00000014847 | Herc6 | 116.848008 | 1.97230496 | 0.45998722 | 4.33509446 | 1.46E-05 | 0.00458115 |
| **IAV vs. Mock** | ENSMAUG00000015073 | | 9.24856206 | 1.46687202 | 0.465342 | 3.50405034 | 0.00045824 | 0.0761195 |
| **IAV vs. Mock** | ENSMAUG00000015796 | Adar | 119.341852 | 1.74603062 | 0.36392156 | 4.7711515 | 1.83E-06 | 0.00072929 |
| **IAV vs. Mock** | ENSMAUG00000016555 | Xdh | 673.123079 | 1.4536058 | 0.42814636 | 3.39863956 | 0.00067722 | 0.09768484 |
| **IAV vs. Mock** | ENSMAUG00000016558 | Ube2l6 | 20.2841472 | 1.6562272 | 0.47425838 | 3.88208358 | 0.00010357 | 0.02223875 |
| **IAV vs. Mock** | ENSMAUG00000016664 | Ddx60 | 103.677813 | 1.77594725 | 0.43444402 | 4.09156205 | 4.28E-05 | 0.01109504 |
| **IAV vs. Mock** | ENSMAUG00000016788 | Gar1 | 53.8052528 | 1.81124174 | 0.47043052 | 3.91953788 | 8.87E-05 | 0.02051014 |
| **IAV vs. Mock** | ENSMAUG00000016905 | | 122.800512 | 2.49289446 | 0.45832613 | 5.46000023 | 4.76E-08 | 3.61E-05 |
| **IAV vs. Mock** | ENSMAUG00000016913 | Mx2 | 1031.78846 | 3.39625503 | 0.45846043 | 7.8502 | 4.15E-15 | 1.83E-11 |
| **IAV vs. Mock** | ENSMAUG00000018005 | | 62.1010941 | 1.46594038 | 0.43516913 | 3.39434831 | 0.00068792 | 0.09768484 |
| **IAV vs. Mock** | ENSMAUG00000018176 | Ccl8 | 148.658814 | 1.66805262 | 0.46913296 | 3.61859539 | 0.00029621 | 0.05113334 |
| **IAV vs. Mock** | ENSMAUG00000018346 | Gbp2b | 46.6527113 | 1.96182149 | 0.47244659 | 4.23477824 | 2.29E-05 | 0.00649728 |
| **IAV vs. Mock** | ENSMAUG00000018871 | Fcgr4 | 33.3494471 | 1.68081568 | 0.47455148 | 3.75684119 | 0.00017207 | 0.0309167 |
| **IAV vs. Mock** | ENSMAUG00000018991 | | 51.523171 | 1.61345546 | 0.46448612 | 3.48010751 | 0.00050121 | 0.07879778 |
| **IAV vs. Mock** | ENSMAUG00000019294 | Hp | 54.3518025 | 1.59254979 | 0.47044832 | 3.46502743 | 0.00053018 | 0.07911325 |
| **IAV vs. Mock** | ENSMAUG00000019385 | Rnf114 | 158.113477 | 1.44786746 | 0.41560323 | 3.465525 | 0.0005292 | 0.07911325 |
| **IAV vs. Mock** | ENSMAUG00000019671 | Gbp7 | 40.0982502 | 1.77586149 | 0.46706397 | 3.79011526 | 0.00015058 | 0.0288192 |
| **IAV vs. Mock** | ENSMAUG00000019884 | Irf7 | 51.8339066 | 2.51972448 | 0.47350521 | 5.6467489 | 1.64E-08 | 1.54E-05 |
| **IAV vs. Mock** | ENSMAUG00000019934 | Dhx58 | 28.09127 | 1.69010452 | 0.47437136 | 3.58628469 | 0.00033542 | 0.05678966 |
| **IAV vs. Mock** | ENSMAUG00000020370 | Irf9 | 68.1029628 | 1.75783614 | 0.42269541 | 4.15389544 | 3.27E-05 | 0.0087203 |
| **IAV vs. Mock** | ENSMAUG00000021773 | Uba7 | 36.0443047 | 2.32691239 | 0.44735307 | 5.2339316 | 1.66E-07 | 9.13E-05 |
| **IAV vs. Mock** | ENSMAUG00000021814 | Ddx58 | 160.376496 | 2.02712186 | 0.42366247 | 4.77051739 | 1.84E-06 | 0.00072929 |
| **IAV vs. Mock** | ENSMAUG00000022167 | Apobec1 | 120.186745 | 2.68053078 | 0.46694968 | 6.08059071 | 1.20E-09 | 2.11E-06 |

**Table S2: Primers**

| Primer | Direction | Sequence |
|---|---|---|
| **SARS-CoV-2 sgN (TRS-L)** | N/A | CTCTTGTAGATCTGTTCTCTAAACGAAC |
| **SARS-CoV-2 sgN (TRS-N)** | N/A | GGTCCACCAAACGTAATGCG |
| **IAV NP** | Forward | GGCGAAGATGCAACAGCAGGTC |
| **IAV NP** | Reverse | GCACCAGACCTTCTGGGAAGTG |
| **ISG15** | Forward | TCTATGAGGTCCGGCTGACA |
| **ISG15** | Reverse | GCACTGGGGCTTTAGGTCAT |
| **MX2** | Forward | AGGCAGTGGTATTGTCACCAG |
| **MX2** | Reverse | ATCACTGATCCCCAGCCCTAT |
| **IRF7** | Forward | ATTTCGGTCGCAGGGATCTG |
| **IRF7** | Reverse | TGCAAGATAAAGCGTCCCGT |
| **CXCL10** | Forward | TCTGAGTGGGACTCAAGGAATC |
| **CXCL10** | Reverse | CCGGTCGGTCATCGATTTTG |
| **CCL5** | Forward | TCTCCACAGCTGTCCTCACT |
| **CCL5** | Reverse | TCCTTCGGGTGACAAAAACGA |
| **AIF-1/IBA-1** | Forward | GGGGAAAAGCCTTTGGACTG |
| **AIF-1/IBA-1** | Reverse | TCAAACTCCATGTACTTCTTCTTGA |
